# CHOP-c-JUN complex plays a critical role in liver proteotoxicity induced by mutant Z alpha-1 antitrypsin

**DOI:** 10.1101/2020.05.04.076752

**Authors:** Sergio Attanasio, Gwladys Gernoux, Rosa Ferriero, Rossella De Cegli, Annamaria Carissimo, Edoardo Nusco, Severo Campione, Jeffrey Teckman, Christian Mueller, Pasquale Piccolo, Nicola Brunetti-Pierri

## Abstract

Alpha-1 antitrypsin (AAT) deficiency is a common genetic disorder with lung and liver involvement. Most patients carry the Z allele in *SERPINA1* that encodes a mutant AAT (ATZ) forming hepatotoxic polymers. We found *CHOP* upregulation and activation in both mouse (PiZ) and human livers expressing ATZ. Compared to controls, juvenile PiZ/*Chop*^*-/-*^ mice showed reduction in hepatic ATZ and transcriptional response to endoplasmic reticulum stress, as consequence of CHOP-mediated increase of *SERPINA1* transcription. CHOP was found to upregulate *SERPINA1* though binding with c-JUN on *SERPINA1* regulatory elements, thus aggravating hepatic accumulation of ATZ. Increased *CHOP* levels were detected in diseased livers of children homozygous for the Z allele.

Compared to adults, AAT deficiency in infants has different severity and prognosis. Based on our findings, CHOP-c-JUN complex upregulates *SERPINA1* transcription and play an important role in the hepatic disease pathogenesis by increasing the burden of proteotoxic ATZ, particularly in the pediatric population.

## INTRODUCTION

Alpha-1 antitrypsin (AAT) encoded by *SERPINA1* gene is an acute phase protein and one of the major circulating proteinase inhibitors mainly synthesized and secreted by hepatocytes. AAT deficiency is a common genetic cause of lung and liver diseases. The vast majority of patients with AAT deficiency (∼95%) are homozygous for the Z mutation in *SERPINA1* gene (PiZZ) that results in a single glutamic acid to lysine substitution at amino acid position 342 (p.Glu342Lys) (1). This mutation changes the protein structure leading to misfolding, polymerization, and accumulation of ATZ in the endoplasmic reticulum (ER) (1). In PiZZ individuals, intrahepatic retention of ATZ results in low circulating levels of AAT and inadequate anti-protease protection in the lower respiratory tract that can lead to progressive lung emphysema (1). PiZZ individuals are also at risk of developing liver disease due to intracellular retention and accumulation of aberrantly folded ATZ leading to hepatitis, cirrhosis, liver failure, and hepatocellular carcinoma (HCC) (2). There is great variability in severity of liver disease among PiZZ homozygotes and a survey of a unique cohort of homozygote individuals identified by an unbiased newborn screening carried out in Sweden showed that about 8% of homozygotes develop clinically significant liver disease in the first four decades of life (3). A greater proportion of this population will likely develop liver injury as they reach older ages. Nevertheless, a number of PiZZ individuals do not manifest clinical symptoms of liver disease throughout their lifetime (4). These data suggest that genetic and/or environmental modifiers play a major role in susceptibility to liver disease. Understanding the mechanisms involved in the pathogenesis of the liver disorder is important to identify disease-modifiers and to develop effective preventive and therapeutic strategies.

Combining results in mouse and human livers expressing ATZ, we identified an important transcriptional regulator of *SERPINA1* expression aggravating the liver disease. For this study, we used transgenic PiZ mice, which have been a valuable model for studying the liver disease induced by ATZ because in the livers of these mice ATZ accumulates within the ER of hepatocytes as Periodic-Acid Shiff-diastase resistant (PAS-D) positive globules, in a nearly identical manner to livers of human patients (5). Because they have been genetically engineered to express the human *SERPINA1* gene harboring the Z mutation that includes the human promoter region (6), they are also useful to investigate *SERPINA1* transcriptional regulation. Importantly, mouse data were validated in human liver tissues from PiZZ patients of various ages with hepatic disease.

## RESULTS

### CHOP is activated in mouse livers expressing ATZ

We previously found up-regulation of genes related to response to ER stress in PiZ livers (7). Among differentially expressed genes, the transcription factor *Chop* was among the most upregulated. Under non-stress conditions, CHOP is typically not expressed at detectable levels and its subcellular location is mainly in cytoplasm whereas stress conditions induce its nuclear translocation (8). Compared to age-matched wild-type controls, *Chop* showed a significant 12-fold upregulation in 6-week-old PiZ mouse livers by real time PCR (**Fig. 1A**). Greater *Chop* upregulation was detected in 6-week-compared to 36-week-old PiZ mouse livers (**Fig. 1A**). Increased expression of CHOP was confirmed at the protein level in 6- and 36-week-old PiZ mouse livers (**Fig. 1B**). Compared to wild-type controls, PiZ mouse livers showed increased CHOP levels by Western blots also on nuclear extracts, although at lower levels compared to mice injected with tunicamycin, an inducer of ER stress and CHOP expression (9) (**Fig. 1C**). Increased nuclear CHOP in PiZ livers was confirmed by immunohistochemistry (**Fig. 1D**) and Gene Set Enrichment Analysis (GSEA) on microarray expression data (7) showed enrichment of up-regulated CHOP target genes compared to wild-type controls (enrichment score [ES]=0.58) (**Fig. 1E**; **Supplementary Fig. S1**). Moreover, by immunohistochemistry, CHOP-positive nuclei were found in hepatocytes, but they were not detected in infiltrates of inflammatory cells (**Supplementary Fig. S2**). In addition, greater *Chop* expression was detected in the parenchymal cell fraction compared to non-parenchymal fraction obtained by liver perfusion of PiZ mice (**Supplementary Fig. S3**).

**Figure 1.**
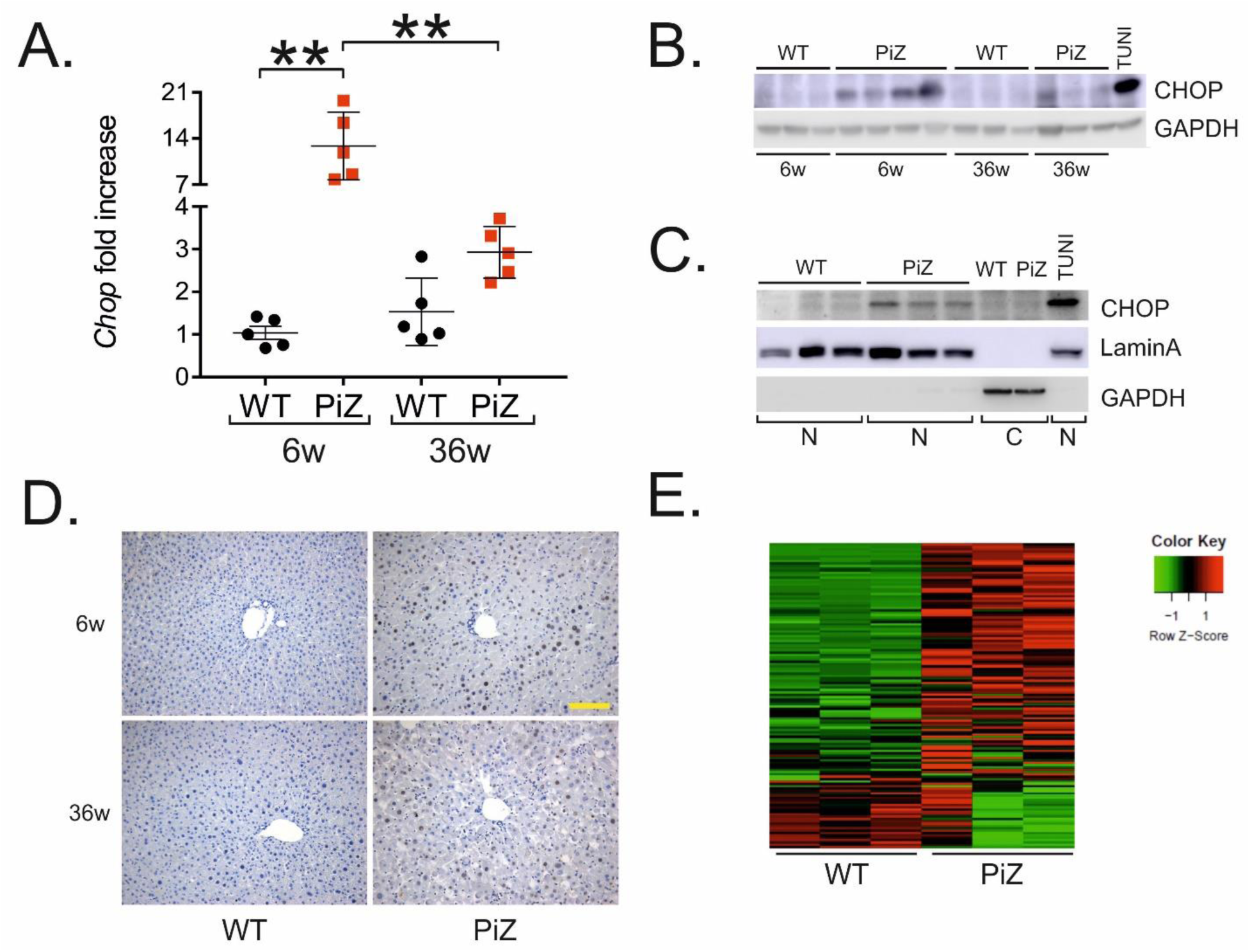
CHOP is activated in PiZ mouse livers. (**A**) *Chop* expression in livers of wild-type (WT) and PiZ mice of 6 and 36 weeks of age (n=5 per group). *β2-Microglobulin* was used for normalization. (**B**) Western blotting for CHOP in livers of WT and PiZ mice of 6 and 36 weeks of age (each lane corresponds to an independent mouse). A liver of a WT mouse injected intra-peritoneally with 4 µg/g tunicamycin (TUNI) was used as a positive control. (**C**) Western blot for CHOP on nuclear extracts of 6-week-old WT and PiZ mouse livers. Lamin A was used as nuclear marker and GAPDH as cytosolic marker. (**D**) Representative immunohistochemistry with anti-CHOP antibody in 6- and 36-week-old WT and PiZ mouse livers (40X magnification; scale bar, 100 µm; n=3 per group). (**E**) Heatmap from GSEA using a set of genes up-regulated by CHOP and differentially expressed genes between 6-week-old PiZ and WT mouse livers. The set of CHOP target genes was defined by combining ChIP-seq and RNA-seq data previously reported from tunicamycin treated wild-type mouse embryonic fibroblasts (42). Averages ± standard deviations are shown. Two-way ANOVA and Tukey’s post-hoc: **p-value < 0.001. *Abbreviations: 6W, 6-week-old; 36W, 36-week-old; N, nuclear extracts; C, cytosolic extracts; TUNI, tunicamycin; WT, wild-type.*

To further investigate CHOP expression and hepatic ATZ accumulation, we injected 4-week-old PiZ mice with a recombinant serotype 8 adeno-associated viral (AAV) vector that incorporates an artificial microRNA (miRNA) targeting the human *SERPINA*1 gene (AAV8-CB-mir914) and results in ATZ hepatic knockdown (10). As controls, mice were injected with the same dose of an AAV8-CB-GFP expressing the green fluorescent protein (GFP). By 4 weeks post-injection mice injected with AAV8-CB-mir914 showed a reduction of hepatic ATZ by PAS-D staining (**Supplementary Fig. S4A**), an approximately 90% reduction of serum ATZ (**Supplementary Fig. S4B**), and about 50% reduction of *Chop* expression compared to mice injected with the control vector (**Supplementary Fig. S4C**). Taken together, these data support upregulation and activation of CHOP in livers of mice expressing ATZ.

### Reduced hepatic ATZ accumulation in *Chop* deleted juvenile PiZ mice

To evaluate the role of CHOP in hepatic ATZ accumulation, we crossed PiZ mice with *Chop*^*- /-*^ mice (9). *Chop*^*-/-*^ mice do not exhibit a substantial phenotype unless they are subjected to stress signals (9). Compared to wild-type, PiZ mice have reduced weight (11) but PiZ/*Chop*^*-/-*^ mice showed greater weight compared to PiZ mice (**Supplementary Fig. S5**). Serum ATZ in PiZ/*Chop*^*-/-*^ mice was lower compared to PiZ mice at 6 weeks of age but no differences were detected in 36-week-old PiZ/*Chop*^*-/-*^ mice (**Fig. 2A**). Compared to PiZ, livers of 6-week-old PiZ/*Chop*^*-/-*^ mice showed marked reduction of hepatic ATZ by PAS-D staining and immunofluorescence with anti-polymer antibody (**Fig. 2B**). However, neither PAS-D staining nor ATZ polymer immunofluorescence detected any difference in ATZ accumulation in livers of 36-week-old PiZ/*Chop*^*-/-*^ mice compared to age-matched PiZ controls (**Fig. 2B; Supplementary Fig. S6**). Consistent with PAS-D staining and ATZ polymer immunofluorescence, compared to age-matched PiZ controls, PiZ/*Chop*^*-/-*^ mice showed reduced ATZ protein by immunoblot performed on soluble and insoluble fractions at 6 weeks of age but not at 36 weeks (**Supplementary Fig. S7**). PiZ/*Chop*^*-/-*^ mouse livers did not show increased apoptosis like PiZ controls, consistent with previous reports (12) (**Supplementary Fig. S8**). Targeted qPCR analysis showed down-regulation of *SERPINA1* expression in PiZ/*Chop*^*-/-*^ compared to age- and gender-matched 6-week-old PiZ mice and consistent with liver ATZ content, no significant differences in *SERPINA1* expression were detected in older mice (**Fig. 2C**). Although CHOP has been involved in liver fibrosis (13), compared to age-matched PiZ mice, liver fibrosis in 69 week-old PiZ/*Chop*^*-/-*^ mice appeared to be unaffected by Sirius red staining and expression of *Col1a1, Timp1*, and *α-Sma* (**Fig. 2D** and **Supplementary Fig. S9A-C**).

**Figure 2.**
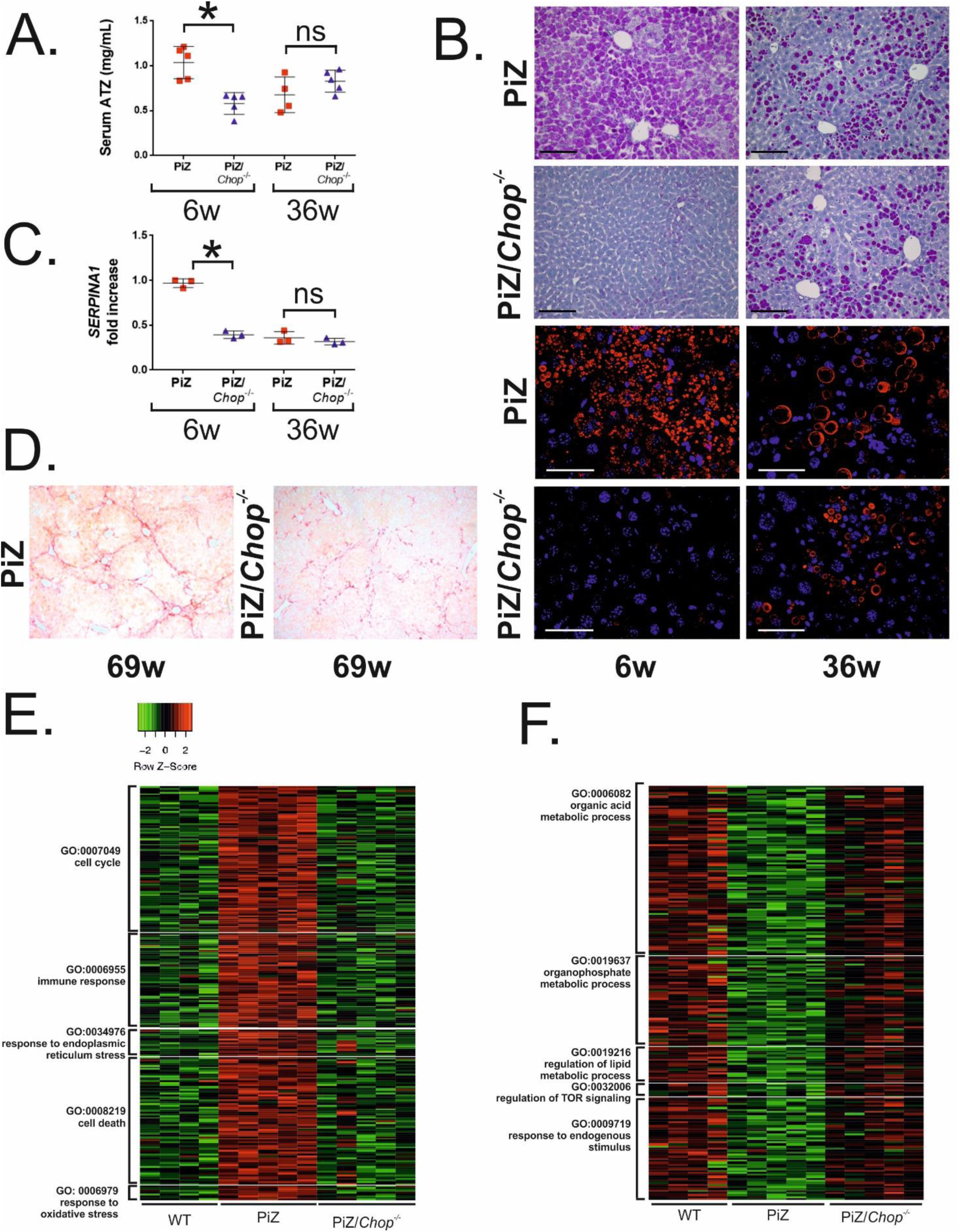
Genetic ablation of *Chop* reduces hepatic ATZ accumulation in juvenile PiZ mice. (**A**) ELISA for serum ATZ in PiZ and PiZ/*Chop*^-/-^ mice of 6 and 36 weeks of age (at least n=4 per group). Serum was collected from the same mice at different ages. (**B**) Representative PAS-D staining (20X magnification, scale bar: 100µm) and immunofluorescence for polymeric ATZ (63X magnifications, scale bar: 50µm) of livers of PiZ and PiZ/*Chop*^-/-^ mice of 6 and 36 weeks of age. (**C**) *SERPINA1* expression in PiZ and PiZ/*Chop*^*-/-*^ mice at 6 and 36 weeks of age (n=3 per group). β2-Microglobulin was used for normalization. Averages ± standard deviations are shown. Two-way ANOVA and Tukey’s post-hoc: *p-value < 0.05, **p-value < 0.005. (**D**) Representative Sirius red staining on livers of 69-week old PiZ and PiZ/*Chop*^*-/-*^ mice (10X magnification). (**E**, **F**) Heatmaps showing the most relevant up- and down-regulated pathways in 6-week-old PiZ mouse livers and normalized to age-matched wild-type levels in PiZ/*Chop*^*-/-*^ mice. *Abbreviations: ns, not statistically significant; 6W, 6-week-old; 36W, 36-week-old; 69W, 69-week-old.*

Unbiased RNA-seq on liver RNA revealed that 6-week-old PiZ/*Chop*^*-/-*^ exhibited an hepatic gene expression profile different from age- and gender-matched PiZ mice and shifting towards wild-type controls on principal component analysis (PCA) (**Supplementary Fig. S10**). In contrast, no obvious differences in gene expression profiles were observed between PiZ/*Chop*^*-/-*^ and PiZ mouse livers of 36-weeks of age, consistent with the ATZ accumulation (**Supplementary Fig. S10**). To specifically investigate the effect of CHOP on gene expression profiles of PiZ mice, we performed a comparison of all datasets, as shown by the VENN diagrams in which we evaluated differentially expressed genes obtained by three different comparisons: PiZ *vs.* wild-type, PiZ/*Chop*^*-/-*^ *vs.* wild-type, and PiZ/*Chop*^*-/-*^ *vs.* PiZ at 6- and 36-weeks of age. This analysis allowed to isolate only genes differentially expressed in PiZ *vs.* wild-type and normalized to wild-type levels in PiZ/*Chop*^*-/-*^ mice. The analysis revealed 912 genes differentially expressed (almost all in opposite correlation) in the intersection between PiZ *vs.* wild-type and PiZ/*Chop*^*-/-*^ *vs.* PiZ (in red in **Supplementary Fig. S11**) whereas at 36 weeks of age, no differences in gene expression profiles between PiZ *vs.* wild-type and PiZ/*Chop*^*-/-*^ *vs.* PiZ mice were detected. Next, we focused only on transcriptomes of 6-week-old mouse livers for further analysis and we found that 467 genes were up-regulated in PiZ mouse livers compared to wild-type controls and normalized to wild-type levels in PiZ/*Chop*^*-/-*^ mice (gene dataset A). In contrast, 444 genes were down-regulated in PiZ mouse livers compared to wild-type controls and normalized to wild-type levels in PiZ/*Chop*^*-/-*^ mice (gene dataset B). Gene ontology (GO) functional analysis was performed by restricting the output to the Biological Processes (BP) in which the two datasets (A and B, separately) were functionally involved. This analysis showed several BP mainly including ER-stress, cell proliferation and immune response for dataset A analysis, and metabolic pathways for dataset B among the most significant (Enrichment score >1.5) (**Supplementary Fig. S12-13**). Next, we dissected clusters relevant for the pathogenesis of liver disease induced by ATZ, including ER-stress, cell cycle, immune response, cell death, and response to oxidative stress that were upregulated in PiZ mouse livers and normalized to wild-type levels in PiZ/*Chop*^*-/-*^ mice (**Fig. 2E**). In contrast, biological clusters of processes related to metabolic functions were downregulated in PiZ mouse livers and normalized to wild-type levels in PiZ/*Chop*^*-/-*^ mice (**Fig. 2F**). Collectively, these data showed that deletion of *Chop* reduces expression of pathways involved ER-stress and immune response while restoring several metabolic functions induced by ATZ accumulation. CHOP is also involved in induction of autophagy genes, such as *Atg*5 and *Atg7* upon nutrients deprivation (14). However, RNA-seq studies did not revealed changes in expression of autophagy genes.

### Complex of CHOP with c-JUN upregulates human *SERPINA1*

Human *SERPINA1* gene contains regulatory elements both at 5’untraslated region (UTR) (**Fig. 3A**) and 3’UTR (15). As previously described, the 5’UTR contains binding sites for AP-1 (in green in **Fig. 3A** (16). CHOP can make stable heterodimers with AP-1 complex (*e.g.*, c-JUN and c-FOS) increasing AP-1mediated gene expression without binding DNA (16). To investigate CHOP-mediated regulation of human *SERPINA1* expression, we co-transfected HeLa cells with a plasmid expressing CHOP and the pAAT-Luc-AAT-3’UTR plasmid that contains the *SERPINA1* regulatory elements included in PiZ transgenic mice (7) (dashed line in **Fig. 3A**) upstream the firefly luciferase coding region and human *SERPINA1* 3’UTR. As positive control, cells were co-transfected with a plasmid expressing c-JUN that transactivates luciferase expression, as previously shown (7). Compared to cells co-transfected with the negative control, cells co-transfected with CHOP showed mild increase in luciferase levels, suggesting that CHOP upregulates *SERPINA1* expression (**Fig. 3B**).

**Figure 3.**
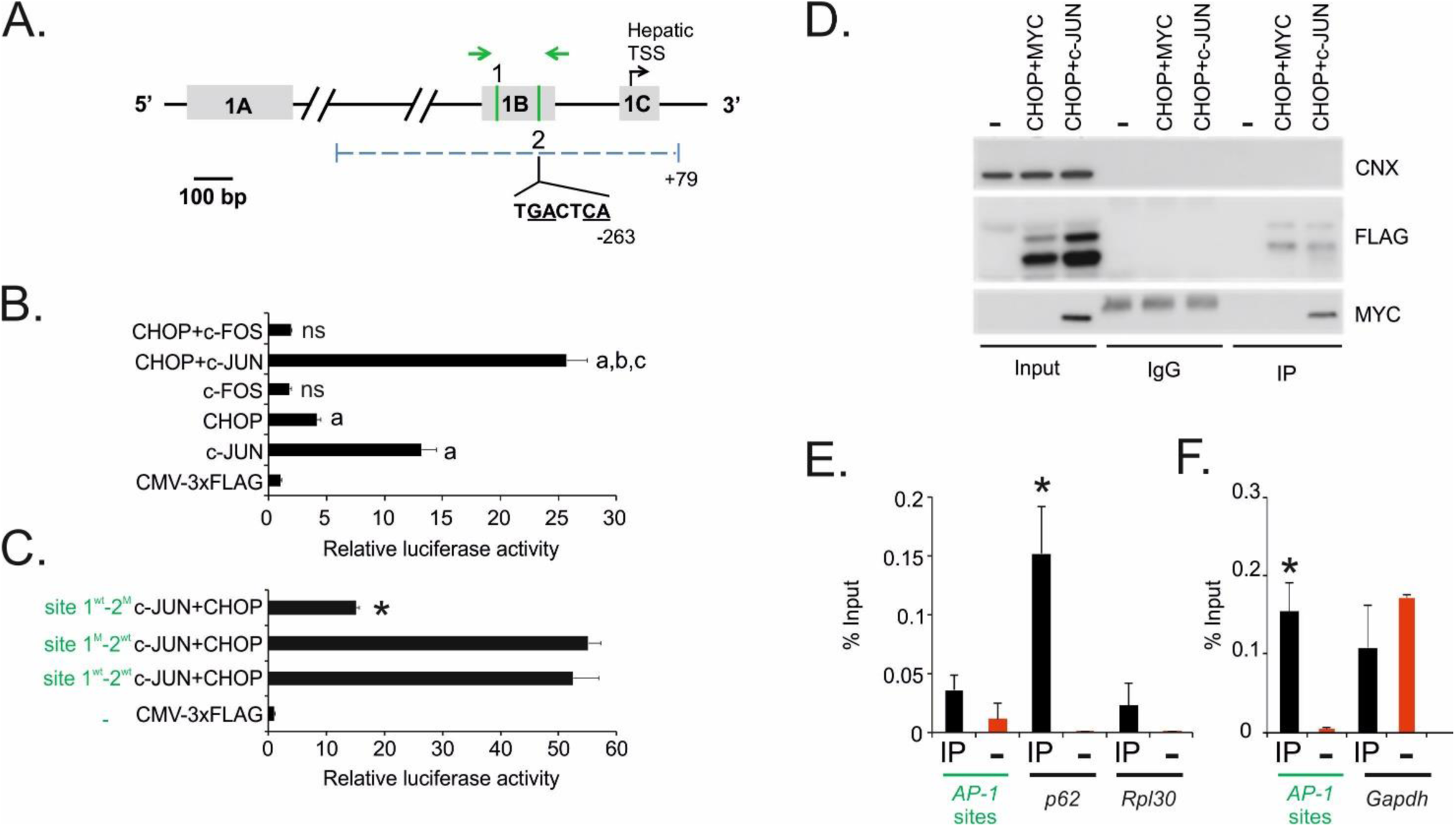
CHOP-c-JUN complex upregulates human *SERPINA1* expression. (**A**) Schematic representation of AP-1 binding sites (1 and 2; in green) in the 5’untraslated region (UTR) of human *SERPINA1* gene (drawn to scale) retained in PiZ mouse genome. Nucleotide sequence is shown for the binding site confirmed by luciferase assays: site 2 for c-JUN-CHOP heterodimer. Nucleotide numbering is calculated as distance from the hepatic TSS. Arrows indicate the position of the primers for the qPCR on the liver chromatin immunoprecipitates shown in E. The dashed line indicates the promoter region cloned upstream the luciferase reporter gene in the pAAT-Luc-AAT-3’UTR plasmid. Underlined nucleotides have been mutagenized. (**B**) Luciferase expression in HeLa cells co-transfected with the pAAT-Luc-AAT.3’UTR construct and the plasmids expressing human CHOP, c-JUN, c-FOS or their combinations. An empty CMV-3XFLAG plasmid was used as control. One-way ANOVA and Tukey’s post-hoc: a, p<0.001 *vs.* CMV-3XFLAG; b, p<0.0001 *vs.* c-JUN; c, p<0.0001 *vs.* CHOP; *ns*, not statistically significant *vs.* CMV-3XFLAG. (**C**) Luciferase expression in HeLa cells co-transfected with the plasmids expressing human CHOP and c-JUN and the pAAT-Luc-AAT.3’UTR construct with mutagenized sites 1 and 2 of CHOP/c-JUN binding sites. An empty CMV-3XFLAG plasmid was used as control. One-way ANOVA and Tukey’s post-hoc: *p-value < 0.0001 *vs.* 1WT-2WT CHOP+c-JUN. (**D**) CHOP and c-JUN co-immunoprecipitation in HeLa cell lines. Cells were transfected with plasmids expressing CHOP-3XFLAG and c-JUN-MYC fusion proteins or control vector (p-CMV-MYC) or left untreated (-). Cell lysates were immunoprecipitated with anti-FLAG antibody and immunoblotted with anti-FLAG antibody to detect CHOP-3XFLAG and anti-MYC antibody to detect c-JUN-MYC. Calnexin (CNX) was used as protein loading control. Anti-FLAG immunoblotting displayed two bands corresponding to the two CHOP isoforms arising from differential translation initiation. (**E**) Chromatin immunoprecipitations (ChIP) on PiZ mouse livers with an anti-CHOP antibody: *p62* and *Rpl30* promoter regions were used as positive and negative controls respectively. (**F**) ChIP on PiZ mouse livers using anti-phospho-c-JUN antibody followed by qPCR on CHOP/c-JUN putative binding sites (*AP-1* sites). *Gapdh* was used as negative control. *t-*test: *p-value < 0.01. *Abbreviations: IP, immunoprecipitates; ns, not statistically significant.*

In contrast, transfections of plasmids expressing c-FOS did not significantly increase luciferase expression (**Fig. 3B**). Co-transfection of CHOP and c-JUN resulted in greater increase of luciferase levels compared to cells transfected with plasmids expressing each of the two transcription factors alone (**Fig. 3B**). Mutagenesis of the binding site 2 but not site 1 of the AP-1 binding site (in green in **Fig. 3A**) reduced luciferase expression after co-transfection of CHOP and c-JUN (**Fig. 3C**). Taken together, these results show that site 2 of the AP1 binding site is required for transactivation by CHOP and c-JUN. Protein co-immunoprecipitation in HeLa cells co-transfected with plasmids expressing FLAG-tagged CHOP and MYC-tagged c-JUN showed that CHOP binds c-JUN (**Fig. 3D**). Real time PCR on chromatin immunoprecipitations (ChIP) of PiZ mouse livers using an anti-CHOP antibody did not detect enrichment for AP-1 binding site sequences (shown in green in **Fig. 3A**) compared to control chromatin without immunoprecipitation (**Fig. 3E**). As positive control of the ChIP with anti-CHOP antibody, *p62* promoter region sequence was enriched in PiZ livers whereas no signal was detected for the *Rpl30* promoter sequence used as negative control. Consistent with previous findings (7), enrichment of AP-1 binding site sequences (shown in green in **Fig. 3A**) was detected in the chromatin immune-precipitates with anti-phospho-c-JUN antibody whereas no enrichment was detected for *Gapdh* promoter region used as negative control (**Fig. 3F**). Altogether, these results suggest that CHOP forms a complex with c-JUN and enhances AP-1 mediated gene expression without direct binding to the DNA.

### Age-dependent upregulation of CHOP in human PiZZ livers

To interrogate the clinical relevance of our findings, we investigated CHOP levels and activation in livers of PiZZ patients. In liver samples from PiZZ patients of less than 16 years of age with extensive PAS-D staining and end-stage liver disease requiring liver transplantation (7) (**supplementary table S1**), increased CHOP nuclear signals was detected compared to controls (**Fig. 4** and **supplementary fig. S14**).

**Figure 4.**
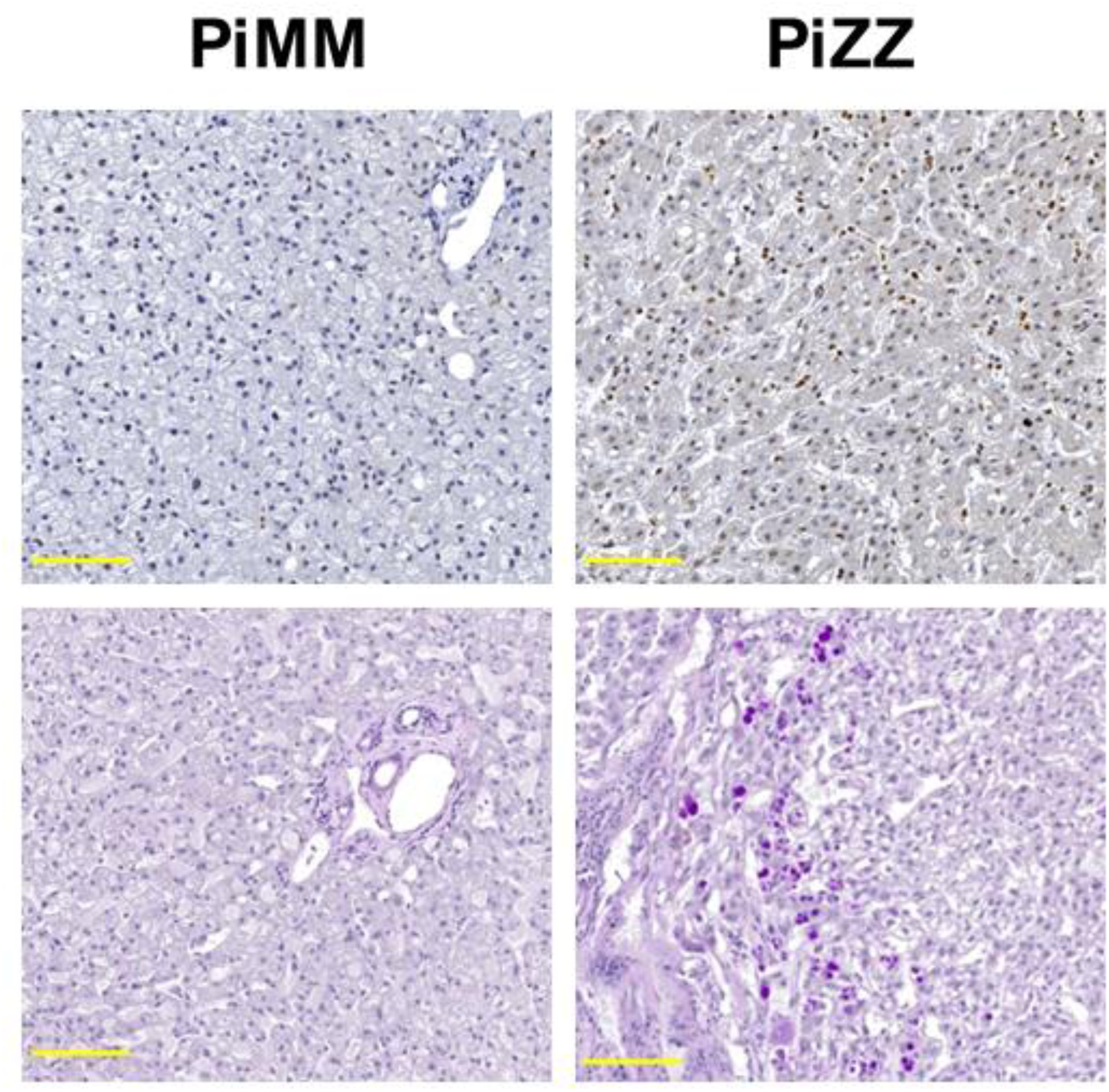
Increased expression and age-dependent upregulation of *CHOP* in human livers expressing ATZ. **(A)** Representative immunohistochemistry with anti-CHOP antibody (upper panels) and corresponding PAS-D staining (lower panels) in liver of a PiZZ patient with end-stage liver disease who underwent liver transplantation (PiZZ #3) and an age-matched PiMM control (PiMM #4) who also underwent liver transplantation as controls (7) (20X magnifications; scale bar, 100µm; the insects are 40X magnifications). Immunohistochemistry with anti-CHOP antibody and PAS-D staining for additional cases are shown in **Supplementary Fig. S4**.

Based on the mouse data, showing a role of CHOP in ATZ accumulation in younger mice but not in older animals, we hypothesized that *CHOP* upregulation is greater in PiZZ children with liver disease compared to PiZZ adults with liver disease. CHOP upregulation was indeed found to be greater in children compared to adults (**Fig. 5** and **supplementary table S2**). Taken together, these results supported a role of CHOP in the pathogenesis of ATZ-related liver disease in children.

**Figure 5.**
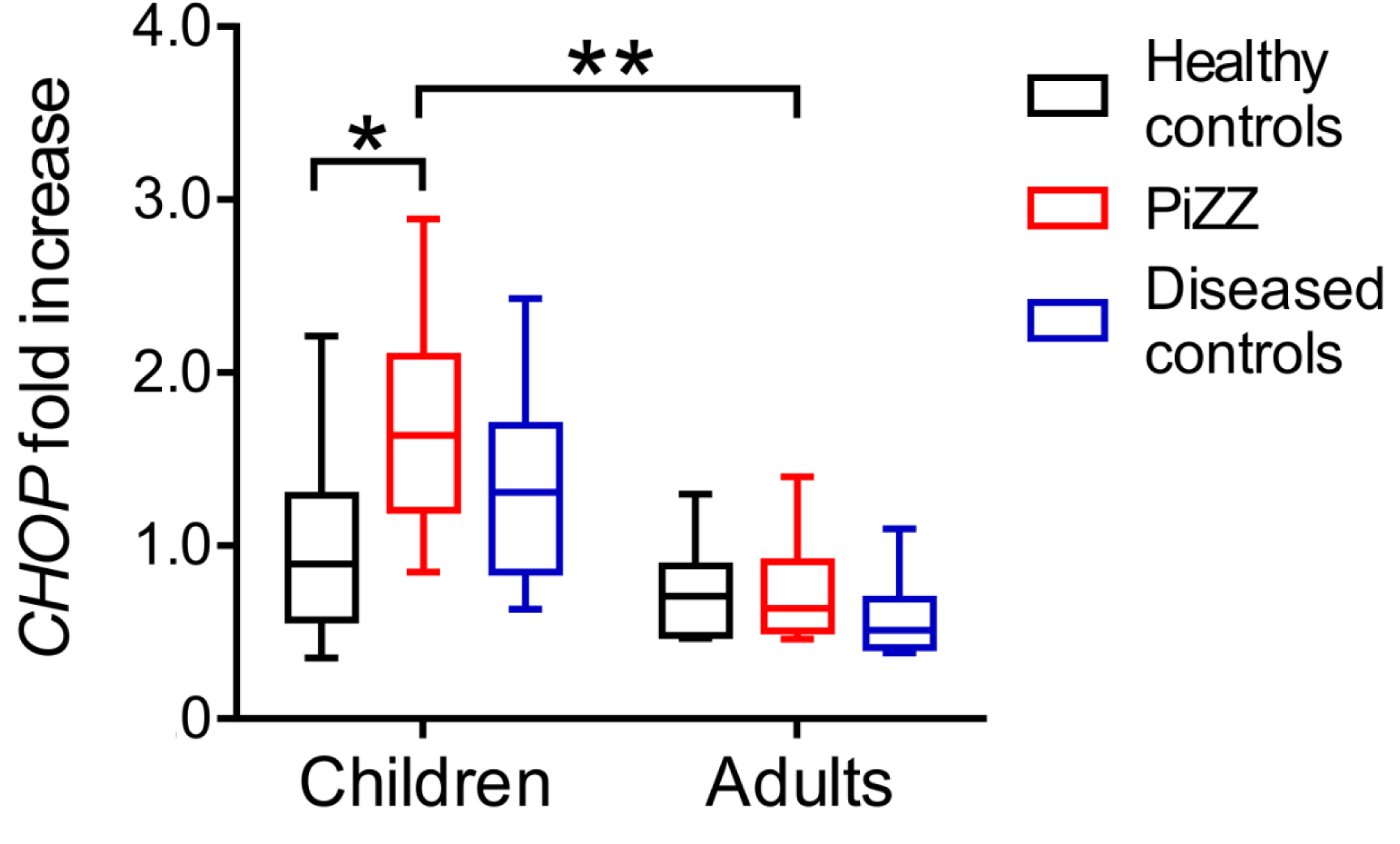
Age-dependent upregulation of *CHOP* in human livers expressing ATZ. ddPCR for *CHOP* expression in human liver biopsies from healthy controls (adults: n=6, age: 50.7±13.3 years; children: n=12, age 6.8±3.6 years), diseased controls (adults: n=6, age: 50.5±13.0 years; children: n=10, age: 5.9±4.3 years), and PiZZ patients (adults: n=6, age: 50.8±13.0 years; children: n=14, age: 6.5±3.7 years). Two-way ANOVA and Tukey’s post-hoc: *p<0.05; **p<0.01.

## DISCUSSION

AAT is an acute phase reactant and its circulating levels may rise several folds in response to several inflammatory stimuli and stress conditions (17). However, in subjects carrying the Z allele, increased synthesis of ATZ augments the burden of mutant protein that correlates with the degree of liver injury and fibrosis (18). To address the factors involved in the pathogenesis of the liver disease, most studies have been focused on ATZ protein degradation by the ubiquitin-proteasome system or autophagy (19) whereas the contribution of transcriptional regulation of *SERPINA1* expression has been neglected for the most part. In this study, we found that CHOP is an important regulator of *SERPINA1* expression, it is upregulated and activated in livers of PiZ mice and PiZZ patients and juvenile *Chop*-deleted mice show marked reduction of liver ATZ accumulation. Moreover, we found an important role of CHOP in regulating c-JUN-mediated upregulation of *SERPINA1*. However, we also found that the lack of CHOP does not affect the progression of the hepatic disease because livers of older mice show ATZ accumulation and liver fibrosis similar to PiZ/*Chop*^*+/+*^ mice. Finally, we detected higher levels of CHOP in children with ATZ-related liver disease compared to adults.

While the precise factors responsible for its activation in livers expressing ATZ remain to be identified, CHOP was found to play an important role in aggravating ATZ accumulation in juvenile PiZ mice through transcriptional upregulation of *SERPINA1* expression. CHOP is an important player in the unfolded protein response and ER-stress response. Moreover, it is activated by several stimuli, such as amino acid and glucose deprivation, DNA damage, cellular growth arrest and hypoxia (20). Previous studies showed that cells expressing ATZ are more sensitive to activate ER stress after a second hit. We also previously found that toxic accumulation of ATZ leads to dysfunctional liver zonation and metabolic alterations (21). Considering these evidences, we speculate that ATZ accumulation may induce such liver alterations including perturbed metabolism that in turn leads to CHOP activation.

We previously found that JNK-mediated activation of c-JUN results in upregulation of *SERPINA1* in human and mouse livers expressing ATZ (7). Here, we showed that CHOP enhances c-JUN-mediated transcriptional upregulation of *SERPINA1.* These findings are consistent with previous studies proposing that CHOP enhances transcriptional activation of AP-1 by tethering to the AP-1 complex without direct binding of DNA (22). Previous data also showed that CHOP and c-JUN also cooperatively mediate the induction of PUMA (p53 upregulated modulator of apoptosis) during hepatocyte lipoapoptosis(23). More broadly, these data raise the question of whether CHOP can potentiate c-JUN-mediated transcription in other cell types under different physiologic and disease conditions. Interestingly, CHOP is transcriptionally positively regulated by the AP-1 complex (24) suggesting a feedback positive loop, in which c-JUN activation upregulates CHOP whereas CHOP activation in turn further enhances *SERPINA1* expression. Such feed-forward mechanism could increase ATZ overload and worsening of the liver disease. Moreover, JNK-mediated activation of c-JUN might also result in long-term consequences on progression of the liver disease that deserve further investigation.

Previous studies showed that CHOP activates GADD34 which dephosphorylates eIF2α and increases protein load (25). Moreover, CHOP was also found to induce oxidative stress which may affect protein folding (26-28). Therefore, in addition to the transcriptional effect, we cannot rule out a beneficial effect of CHOP deletion on improved protein folding by reduction of protein load and oxidative stress. By immunohistochemistry and analysis of parenchymal and non-parenchymal liver fractions, *Chop* expression was detected largely in hepatocytes. However, additional effects of CHOP in other cell types besides hepatocytes cannot be ruled out.

In contrast to older mice, juvenile PiZ mice deleted for *Chop* showed marked reduction of ATZ accumulation suggesting a different role of CHOP at different disease stages. Interestingly, levels of blood ATZ in PiZZ patients appear to decline over time (29), a finding that parallels the mouse data shown in this and previous studies (17). The mechanisms underlying the age dependent reduction of CHOP in livers expressing ATZ are unknown but the reduced proteotoxic load due to recently recognized decreased of HNF-4α is a possible explanation (21).

Clinical data suggest that the human disease has different features in infants compared to adults. A survey of large transplantation databases found that AAT deficiency most commonly causes liver disease requiring liver transplantation in adults, but it also revealed that almost all transplants in children were performed under the age of 5 years (30). These differences suggest that pathogenetic mechanisms vary depending on the age and more potent modifiers play a role in the liver disease in children (30). Based on our findings, we propose that CHOP and factors regulating its expression might be among the modifiers responsible for the more severe disease occurring in the pediatric population. Interestingly, CHOP is upregulated by non-steroidal anti-inflammatory drugs (NSAIDs) (31) and indomethacin given to PiZ mice was previously shown to increase hepatic injury and *SERPINA1* expression resulting in greater accumulation of ATZ (32). Moreover, exposure of hepatocytes to acetaminophen activates JNK and CHOP and disruption of Jnk2 or *Chop* reduce acetaminophen-induced liver injury (33). Therefore, environmental factors, including drugs, could aggravate ATZ-induced liver injury and may be dangerous particularly in children with AAT deficiency.

In conclusion, our study suggest that CHOP is an important player in aggravating hepatic accumulation of toxic ATZ in mice and in patients, especially at younger ages. Drugs inhibiting CHOP activity might have potential for therapy of liver disease in PiZZ children.

## MATERIALS AND METHODS

### Mouse studies

PiZ transgenic mice (6) were maintained on a C57BL/6 background. *Chop*^-/-^ mice were purchased from Jackson laboratory and bred with PiZ mice. As controls, C57BL/6 (Charles River Laboratories) were treated with DMSO (Sigma) or 4µg/g of Tunicamycin (Sigma). Only male mice were used for all experiments. Blood samples were collected by retro-orbital bleedings. The rAAV8pCB-mir914-GFP and rAAV8pCB-GFP were generated, purified, and titered by the UMass Gene Therapy Vector Core as previously described (10). Vectors were diluted in saline solution and intravenously injected in 4-week-old PiZ mice at the dose of 1×10^13^ genome copies (gc)/Kg.

Liver protein extracts and sections, and serum samples were analyzed by PAS-D staining, immunofluorescence for polymeric ATZ, and ELISA for total human AAT, respectively as described previously (34, 35). Stained liver sections were examined under a Zeiss LSM700 for fluorescence and Leica DM5000 microscope for bright field. Nuclear and cytoplasmic protein extracts were prepared using NXTRACT kit (Sigma-Aldrich) according manufacturer’s instructions.

For real time PCR, total RNA was extracted using RNeasy kit (Qiagen) and 1µg of RNA was reverse transcribed using High Capacity cDNA Reverse Transcription kit according to manufacturer’s protocol (Applied Biosystems). PCR reactions were performed using SYBR Green Master Mix (Roche). PCR conditions were as follows: pre-heating, 5 min at 95°C; cycling, 40 cycles of 15 sec at 95°C, 15 sec at 60°C and 25 sec at 72°C. Results were expressed in terms of cycle threshold (Ct). Ct values were averaged for each duplicate. *β2-microglobulin* (*B2m*) was used as endogenous controls. Data were analyzed using LightCycler 480 software, version 1.5 (Roche). Primer sequences are listed in **supplementary table S3**.

For Western blotting, liver samples homogenized in RIPA buffer with protease and phosphatases inhibitors cocktail (Roche) were incubated for 20min at 4°C and centrifuged at 13,200 rpm for 10min. 10-20µg of lysates were electrophoresed on a 12% sodium dodecyl sulfate (SDS)- PAGE. After transfer to nitrocellulose membrane, blots were blocked in TBS-Tween20 with 5% non-fat milk for 1 hour at room temperature, and the primary antibody was applied overnight at 4°C. Anti-rabbit IgG or anti-mouse IgG conjugated with horseradish peroxidase (GE Healthcare, 1:3,000) and ECL (Pierce) were used for detection. Primary antibodies are listed in **supplementary table S4**.

For immunohistochemistry (IHC), 5µm-thick sections were rehydrated and permeabilized in phosphate-buffered saline (PBS)/0.5% Triton (Sigma) for 20min. Antigen unmasking was performed in 0.01M citrate buffer in a microwave oven. Next, sections underwent blocking of endogenous peroxidase activity in methanol/1.5% H2O2 (Sigma) for 30min and were incubated with blocking solution [3% bovine serum albumin (Sigma), 5% donkey serum (Millipore), 1.5% horse serum (Vector Laboratories) 20mM MgCl2, 0.3% Triton (Sigma) in PBS] for 1 hour. Sections were incubated with primary antibody (**supplementary table S4**) overnight at 4°C and with universal biotinylated horse anti-mouse/rabbit immunoglobulin G secondary antibody (Vector Laboratories) for 1 hour. Biotin/avidin-horseradish peroxidase signal amplification was achieved using the ABC Elite Kit (Vector Laboratories) according to manufacturer’s instructions. 3,30-Diaminobenzidine (Vector Laboratories) was used as peroxidase substrate. Mayer’s hematoxylin (Bio-Optica) was used for counterstaining. Sections were dehydrated and mounted in Vectashield (Vector Laboratories). Image capture was performed using the Leica DM5000 microscope.

### QuantSeq 3’ mRNA sequencing

Preparation of libraries was performed with a total of 100ng of RNA from each sample using QuantSeq 3’mRNA-Seq Library prep kit (Lexogen, Vienna, Austria) according to manufacturer’s instructions. Total RNA was quantified using the Qubit 2.0 fluorimetric Assay (Thermo Fisher Scientific). Libraries were prepared from 100ng of total RNA using the QuantSeq 3’ mRNA-Seq Library Prep Kit FWD for Illumina (Lexogen GmbH). Quality of libraries was assessed by screen tape High sensitivity DNA D1000 (Agilent Technologies). Libraries were sequenced on a NovaSeq 6000 sequencing system using an S1, 100 cycles flow cell (Illumina Inc.). Amplified fragmented cDNA of 300 bp in size were sequenced in single-end mode with a read length of 100 bp. Illumina novaSeq base call (BCL) files are converted in fastq file through bcl2fastq [http://emea.support.illumina.com/content/dam/illuminasupport/documents/documentation/software_documentation/bcl2fastq/bcl2fastq2-v2-20-software-guide-15051736-03.pdf] (version v2.20.0.422). For analysis, sequence reads were trimmed using bbduk software (https://jgi.doe.gov/data-and-tools/bbtools/bb-tools-user-guide/usage-guide/) (bbmap suite 37.31) to remove adapter sequences, poly-A tails and low-quality end bases (regions with average quality below 6). Alignment was performed with STAR 2.6.0a3 (36) on mm10 reference assembly obtained from cellRanger website (https://support.10xgenomics.com/single-cell-gene-expression/software/release-notes/build#mm10_3.0.0; Ensembl assembly release 93). Expression levels of genes were determined with htseq-count (37) using Gencode/Ensembl gene model. We filtered out all genes having <1cpm in less than n_min samples and Perc MM reads >20% simultaneously. Differential expression analysis was performed using edgeR (38), a statistical package based on generalized linear models, suitable for multifactorial experiments. Gene Ontology and Functional Annotation Clustering analyses were performed using DAVID Bioinformatic Resources (39, 40). Data were deposited in GEO with the accession number GSE141593.

### Gene Set Enrichment Analysis

Gene Set Enrichment Analysis (GSEA) (41) was performed by GSEA software (www.broadinstitute.org/gsea). Expression microarray of livers from PiZ mice and wild-type controls was obtained from previous studies (GSE93115), CHOP target genes set was defined by combining ChIP-seq and RNA-seq data previously reported from tunicamycin treated wild-type mouse embryonic fibroblasts (42).

### Protein immunoprecipitation

The pCMV-c-JUN-MYC plasmid was generated by cloning the human c-JUN coding sequence in the pCMV-MYC plasmid (ThermoFisher). Primers used for generation of the construct are shown in **supplementary table S5**. HeLa cells were co-transfected with plasmids expressing CHOP-FLAG and c-JUN-MYC. Cells untreated cells or co-transfected with the plasmid expressing CHOP-FLAG were used as controls. 48 hours after transfection, cells were washed with PBS and then, protein crosslinking was performed using a solution of 1.5mM dithiobis(succinimidylpropionate (DSP) (Thermo Fisher Scientific) in PBS added to the dishes. DSP is a bifunctional crosslinking agent containing an amine-reactive N-hydroxysuccinimide esters that react with primary amines to form stable amide bonds and is more efficient than formaldehyde as protein-protein cross-linker (43). Dishes were incubated with DSP for 2 hours at 4°C and the reaction was stopped with addition for 15min at room temperature of 1M Tris, pH 7.8, to a final concentration of 20mM. Dishes were washed with PBS and then lysed as described above. 1mg of protein lysates was used for the immunoprecipitation with mouse anti-FLAG antibody or mouse IgG overnight at 4°C. The day after, 20µL of Dynabeads Protein G (Thermo Fisher Scientific) were added and incubated for 2 hours at 4°C. The flow through was collected using the magnet and the complex Dynabeads-Ab-Ag was washed 5 times using RIPA buffer with 0.1% Triton without SDS and once with RIPA buffer without Triton and SDS. Finally, the elution was performed by boiling tubes at 100°C for 5 minutes in a solution containing SDS 8%. Samples were analyzed by SDS-PAGE as described above. Primary antibodies used for immunoprecipitations are listed in **supplementary table S4**.

### Luciferase assays

The pAAT-Luc-AAT.3’UTR plasmid was generated by cloning *SERPINA1* 3’untranslated region (UTR) downstream the firefly luciferase coding region in the previously generated pAAT-Luc vector. The cDNA of human CHOP and c-FOS were amplified by RT-PCR from Huh7 cells and cloned into p3xFLAG-CMV-14 vector. The plasmid expressing human c-JUN was previously generated (35). AP-1 sites in the pAAT-Luc-AAT.3’UTR plasmid were mutagenized as previously described (35). Primers used for generation of the constructs are shown in **supplementary table S5**. HeLa cells were cultured in DMEM plus 10% fetal bovine serum (FBS) and 5% penicillin/streptomycin. Cells were co-transfected with the plasmid containing the wild-type or mutagenized AP-1 consensus sequences and with a plasmid expressing the human CHOP, and c-JUN. Each well was co-transfected with the pRL-TK plasmid (Promega) expressing the renilla luciferase as control. Lipo D293 (SigmaGen Laboratories) was used for the DNA transfection according to the manufacturer’s instructions. Cells were harvested 48 hours after transfection and assayed for luciferase activity using the Dual-Luciferase Reporter (DLR™) Assay System (Promega). Data were expressed relative to renilla luciferase activity to normalize for transfection efficiency. Transfections were repeated at least three times.

### Chromatin immunoprecipitations

Chromatin immunoprecipitation (ChIP) was performed as previously described (35). Briefly, PiZ mouse livers were mechanically homogenized in PBS in the presence of protein inhibitors and then crosslinked by formaldehyde to a final concentration of 1% for 10min at room temperature. The cross-linking reaction was stopped by the addition of glycine at a final concentration of 0.125mM for 5min at room temperature. After three washes in PBS with protein inhibitors, liver homogenates were lysed in cell lysis buffer (piperazine-1,4-bis-2-ethanesulfonic acid 5 mM [pH 8.0], Igepal 0.5%, KCl 85 mM) for 15min. Nuclei were lysed in lysis buffer (Tris HCl [pH 8.0] 50mM, ethylene diamine tetra-acetic acid 10mM, sodium dodecyl sulfate 0.8%) for 30min. Chromatin was sonicated to yield DNA fragments approximately 200-1,000 bp. DNA was co-immunoprecipitated using the ChIP-grade anti-phospho-c-JUN (Ser73), and anti-CHOP antibodies overnight at 4°C. No antibody was used in the negative control of the immunoprecipitation. Purified immunoprecipitated DNA samples and inputs were amplified by quantitative PCR with primers specific for the AP-1 binding sites of the 5’ untranslated region (5’-UTR) of the *SERPINA1* gene. Antibodies for ChIP and primer sequences are listed in **supplementary tables S4** and **S6**, respectively.

### Human liver samples

Liver samples for immunohistochemistry were collected anonymously from subjects with end-stage liver disease homozygous for the Z or M alleles of *SERPINA1*. Diseased liver controls were obtained from patients with diagnoses of biliary atresia or cirrhosis. Frozen liver powdered samples were homogenized in Trizol (Invitrogen) and total RNA was extracted according to the manufacturer’s recommendations. 500ng of RNA were retrotranscribed using High capacity RNA to cDNA kit (Applied biosystems) and 2µl were used as an input for droplet digital PCR (ddPCR). For gene expression, *DDIT3* (*CHOP*) gene (unique assay ID: dHasEG5002009, Biorad) was used as a target and *POLR2H* gene (unique assay ID: dHasEG5005571, Biorad) as a reference. ddPCR was run according to PrimePCR ddPCR Gene expression assays (EvaGreen®) protocol (Biorad).

### Statistical analyses

Data are expressed as averages ± standard deviation. Two tailed Student *t* test and analysis of variance (ANOVA) plus Tukey’s post hoc analysis were used as statistical tests for mean comparisons.

### Study approvals

Mouse procedures were performed in accordance to regulations of the Italian Ministry of Health and were conformed to the Guide for the Care and Use of Laboratory Animals published by the National Institutes of Health (NIH publication 86-23, revised 1985). Human liver samples were obtained from the St. Louis University School of Medicine, (St. Louis, MO, USA) and UMass (Boston, MA, USA). Ethics Committees at each institution approved the study. Written informed consent was received from participants prior to inclusion in the study.

## AUTHOR CONTRIBUTIONS

SA performed study concept and design, acquisition of data, analysis and interpretation of data; GG and CM acquired data in human samples; AC performed statistical analyses; EN provided technical support; SC analyzed liver samples; JT provided human samples; PP performed study concept and design, analysis and interpretation of data; NB-P performed study concept and design, analysis and interpretation of data, and wrote the manuscript.

## ACKNOWLEDGEMENTS

This work was supported by grants of Fondazione Telethon Italy to N.B.-P. and Alpha-1 Foundation (Research grant to N.B.P. and Gordon L. Snider Award 2016 to P.P.) and the “Federico II” University of Naples (STAR Program) to P.P. SA is supported by an ALTA award.

## Supplementary Figures and Tables

CHOP-c-JUN complex plays a critical role in liver proteotoxicity by mutant Z alpha-1 antitrypsin.

*Sergio Attanasio, Gwladys Gernoux, Rosa Ferriero, Rossella De Cegli, Annamaria Carissimo, Edoardo Nusco, Severo Campione, Jeffrey Teckman, Christian Mueller, Pasquale Piccolo, and Nicola Brunetti-Pierri*

**Supplementary Figure S1:**
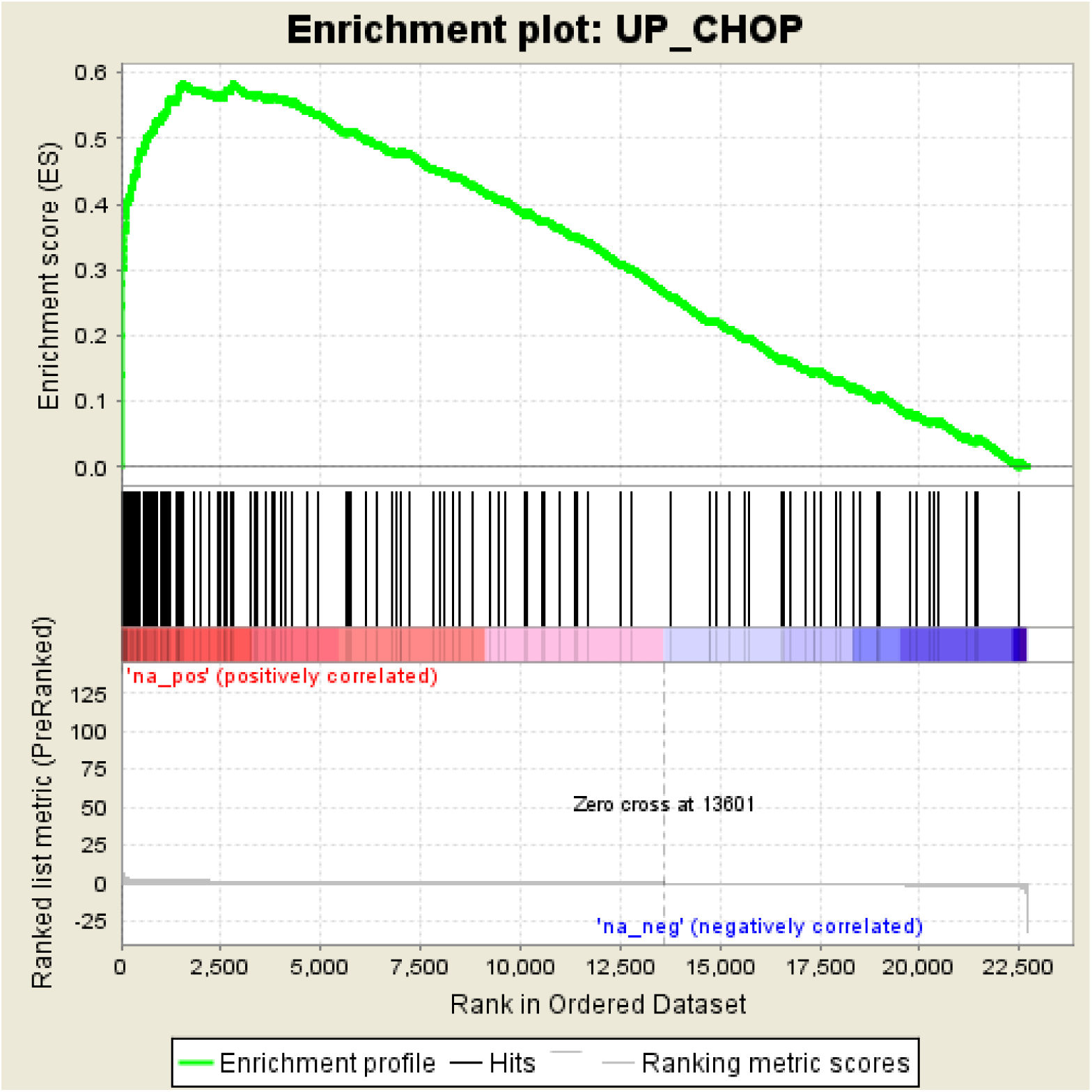
Enrichment plot from Gene Set Enrichment Analysis (GSEA) of genes up-regulated specifically by CHOP shows enrichment in PiZ versus wild-type (WT) mouse livers.

**Supplementary Figure S2:**
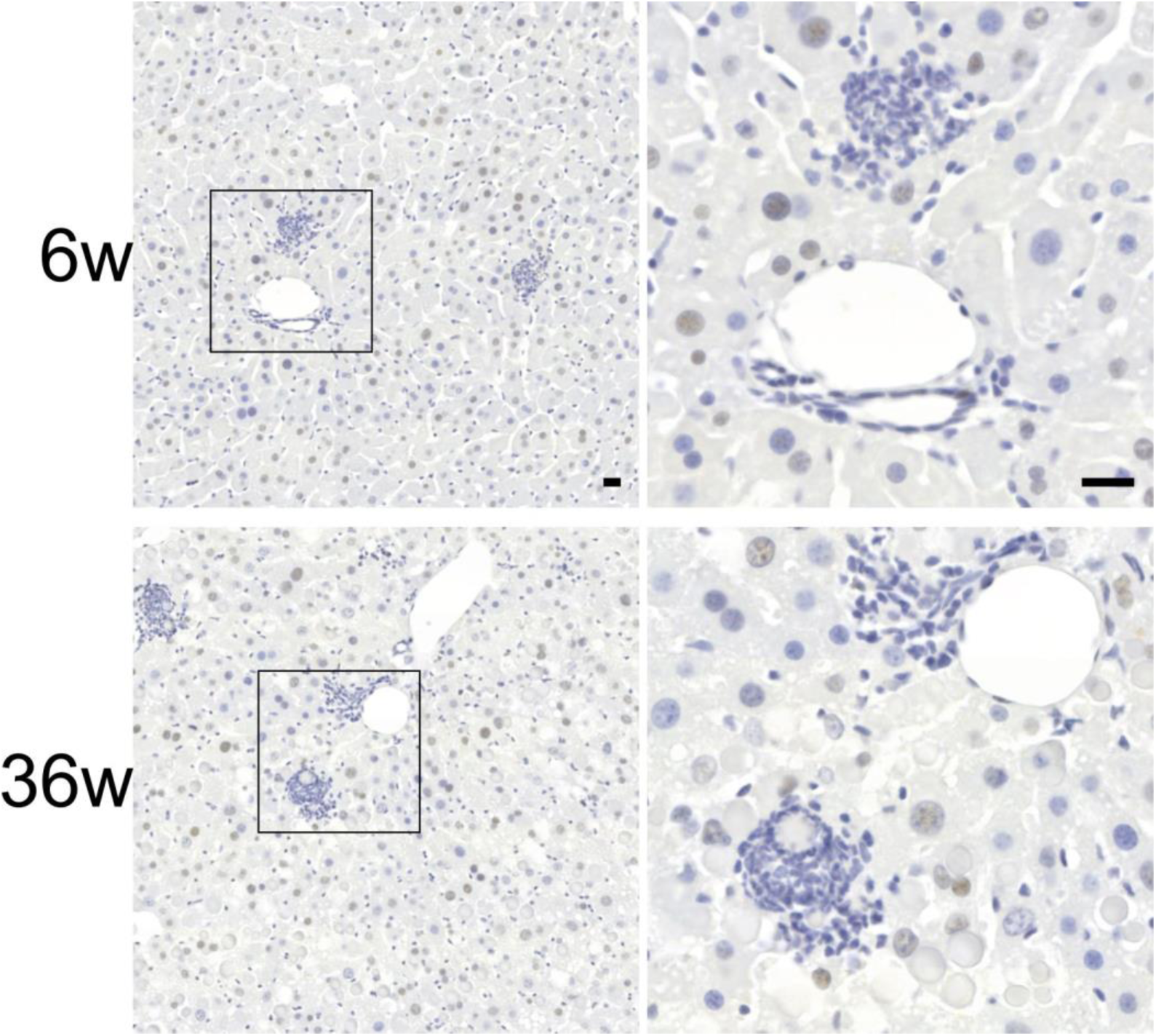
CHOP expression detected in nuclei of hepatocytes but not of inflammatory infiltrating cells. Representative immunohistochemistry with anti-CHOP antibody in livers of PiZ mice of 6 and 36 weeks of age. Images on the right correspond to the insets of left panels. Scale bar: 20µm. *Abbreviations: 6w, 6 weeks; 36w, 36-weeks of age.*

**Supplementary Figure S3:**
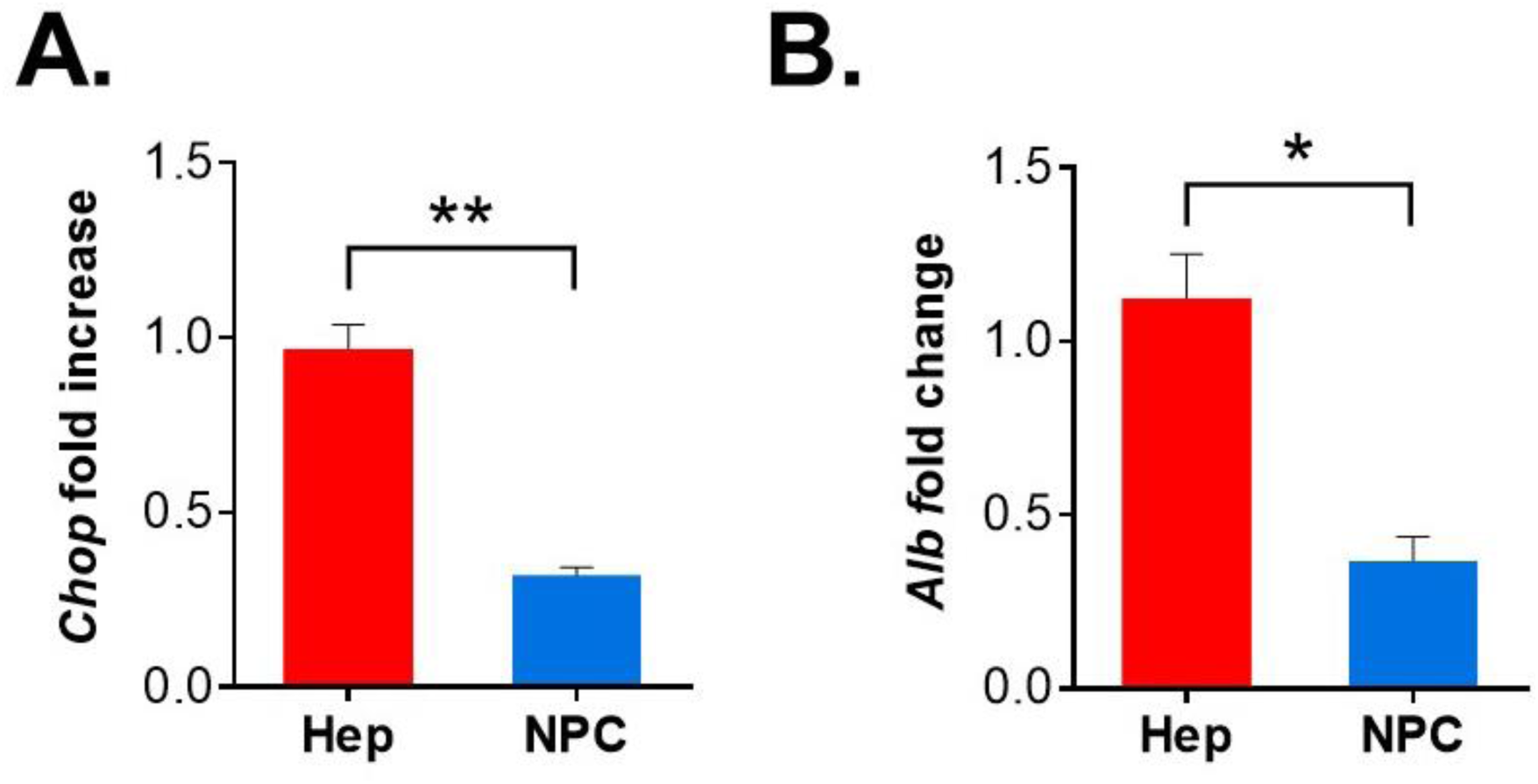
*Chop* expression in parenchymal and non-parenchymal cell enriched fractions of PiZ livers. *Chop* expression is greater in the parenchymal fraction enriched for hepatocytes (Hep) compared to the non-parenchymal cells (NPC). Albumin (*Alb*) gene expression was evaluated to confirm enrichment for hepatocytes. *p<0.05; **p<0.01.

**Supplementary Figure S4:**
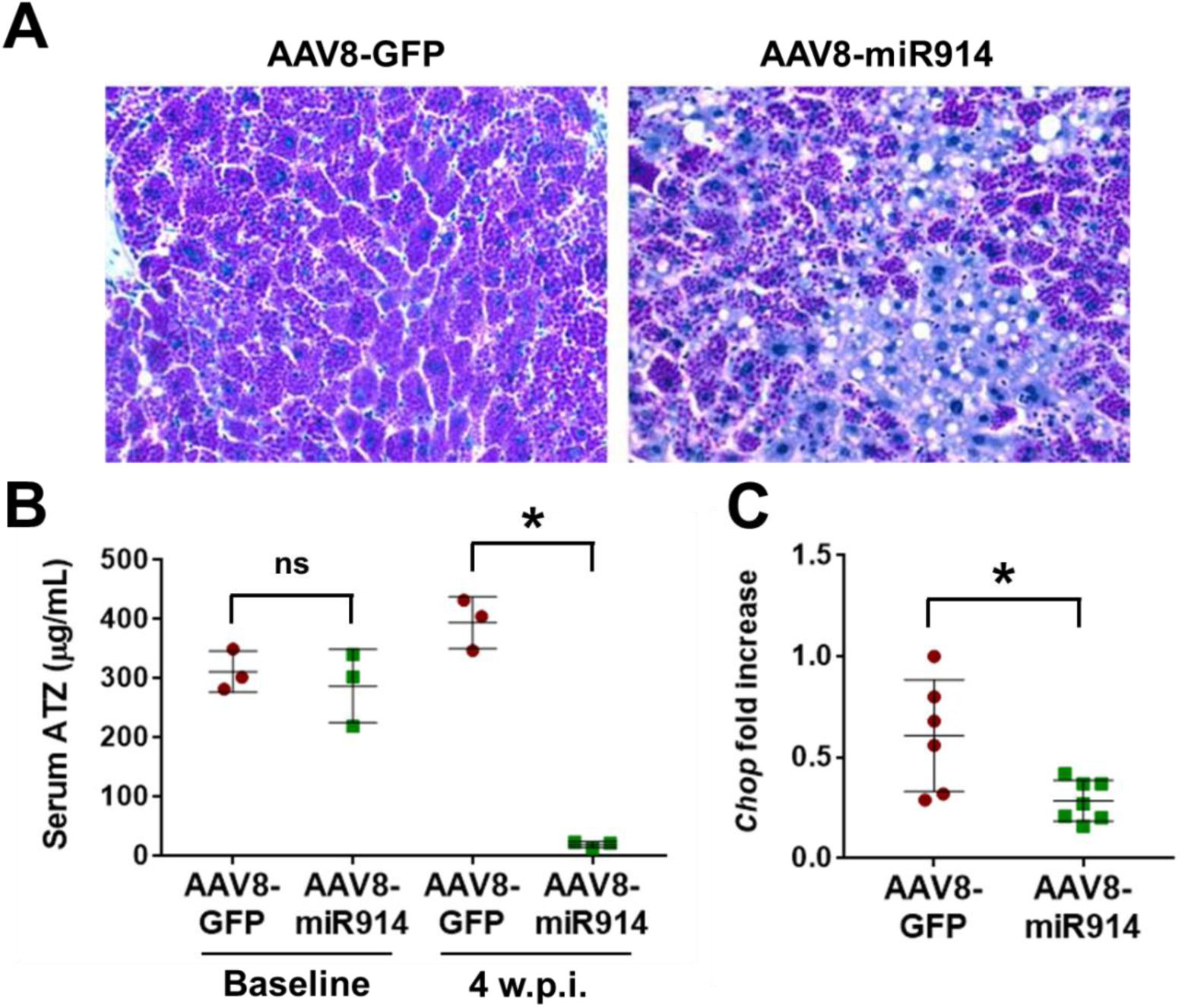
(**A**) Representative PAS-D staining on livers of PiZ mice injected with 1×10^13^ genome copies (gc)/Kg of recombinant serotype 8 adeno-associated viral (rAAV) vector that incorporates microRNA (miRNA) sequences targeting *SERPINA1* gene (AAV8-CB-mir914) or green fluorescent protein (GFP) as control (AAV8-CB-GFP) (20X magnification, at least n=5 per group). (**B**) ELISA for ATZ on serum of PiZ mice at the baseline and 4-weeks after the injection of AAV8-CB-mir914 or AAV8-CB-GFP vectors. (**C**) Real time PCR for *Chop* in livers of PiZ mice injected with AAV8-CB-mir914 or AAV8-CB-GFP vectors (at least n=5 per group). Averages ± SD are shown. *t-*test: *p-value < 0.05. *Abbreviations: ns, not statistically significant; w.p.i., weeks post-injection.*

**Supplementary Figure S5:**
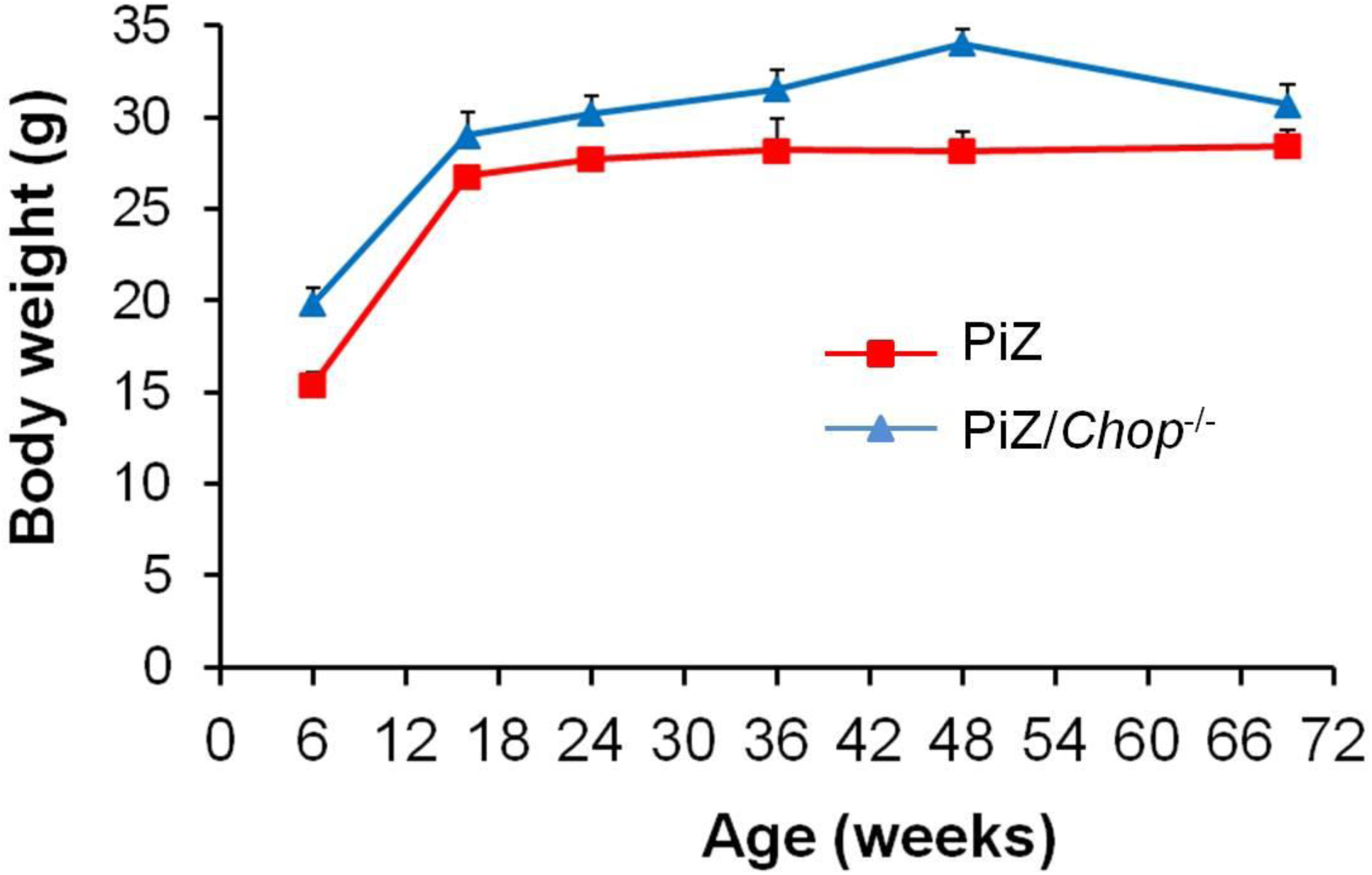
Body weight of PiZ and PiZ/*Chop*^-/-^ mice at different ages (at least n=5 per group). The differences of body weights between PiZ and PiZ/*Chop*^-/-^ mice were statistically significant (Time series analysis; Bayesian factor: 5.478). Averages ± SEM are shown.

**Supplementary Figure S6:**
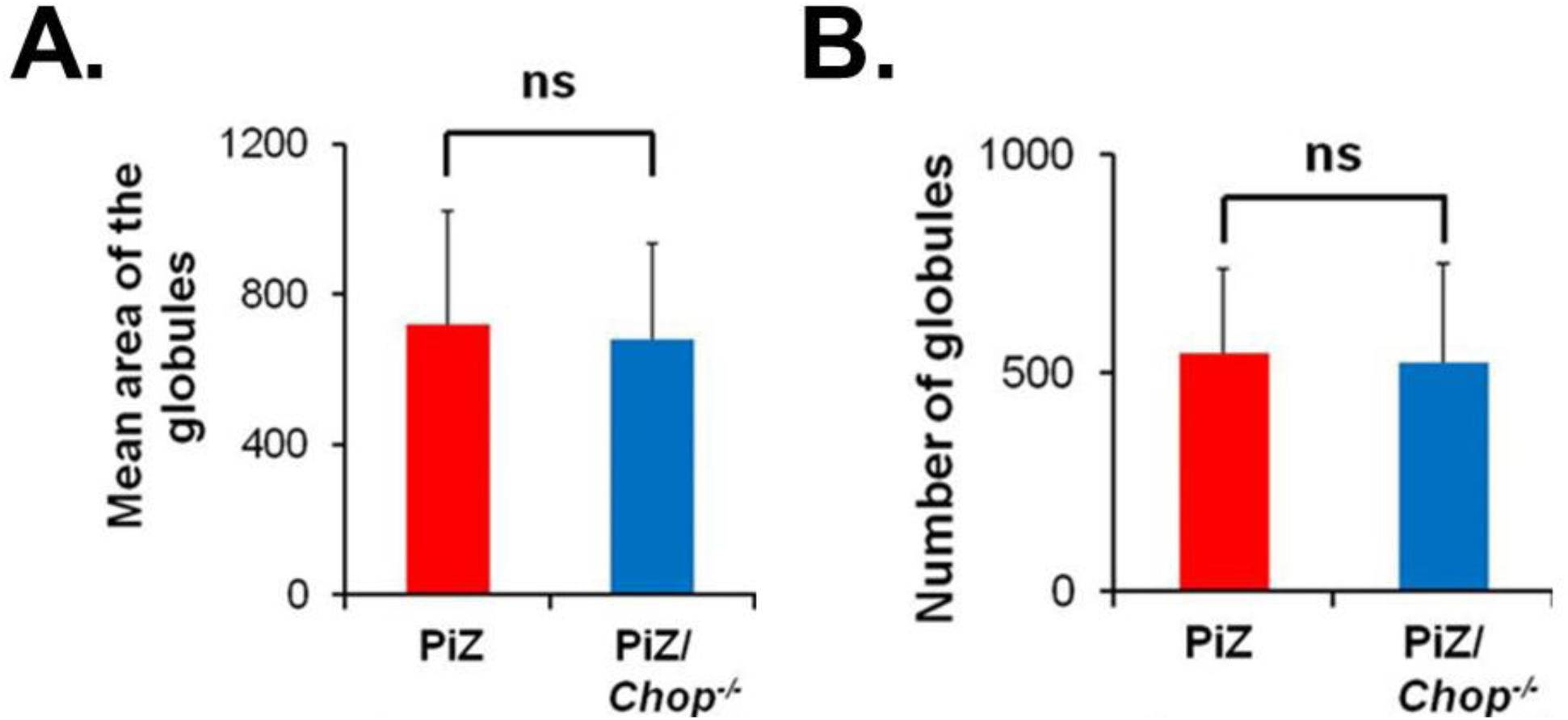
Quantification of the mean area (**A**) and number (**B**) of ATZ globules in 36-week-old PiZ and PiZ/*Chop*^*-/-*^ mouse livers visualized by PAS-D staining (n=3 per group). *Abbreviations: ns, not statistically significant.*

**Supplementary Figure S7:**
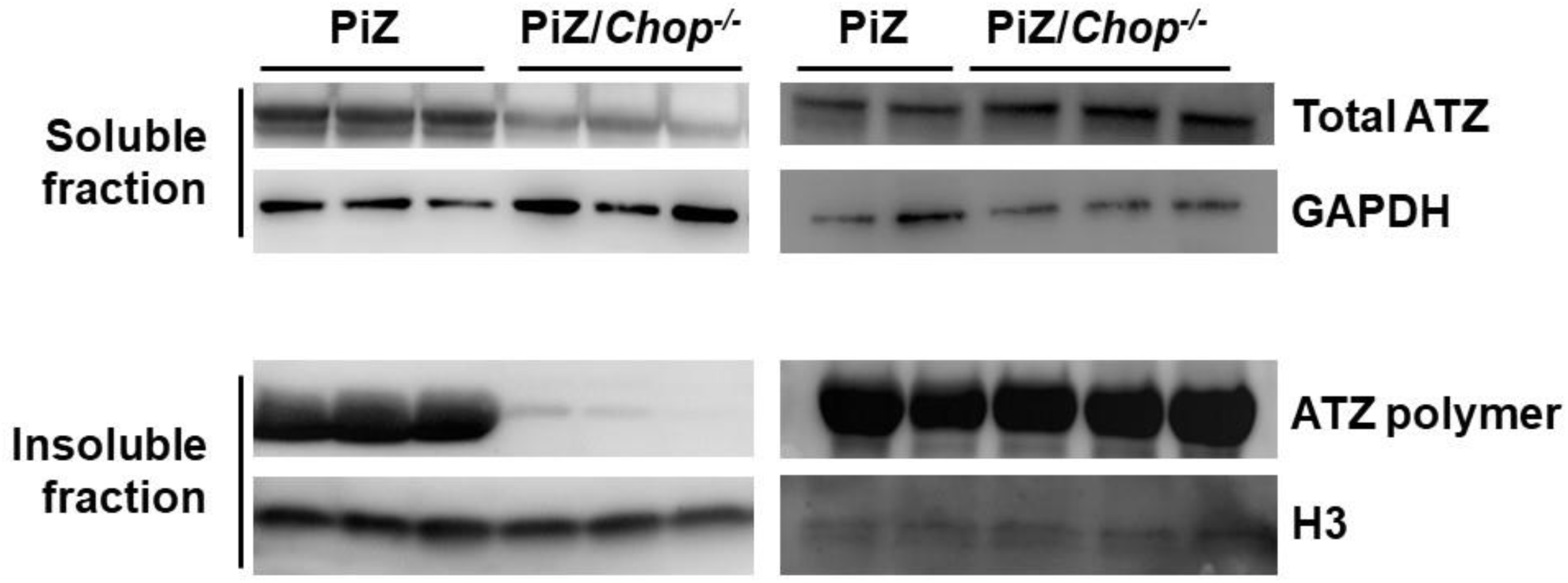
Western blot on soluble and insoluble fractions of 6-week and 36-week old PiZ and PiZ/*Chop*^*-/-*^ mouse livers. GAPDH and H3 were used for normalization.

**Supplementary Figure S8:**
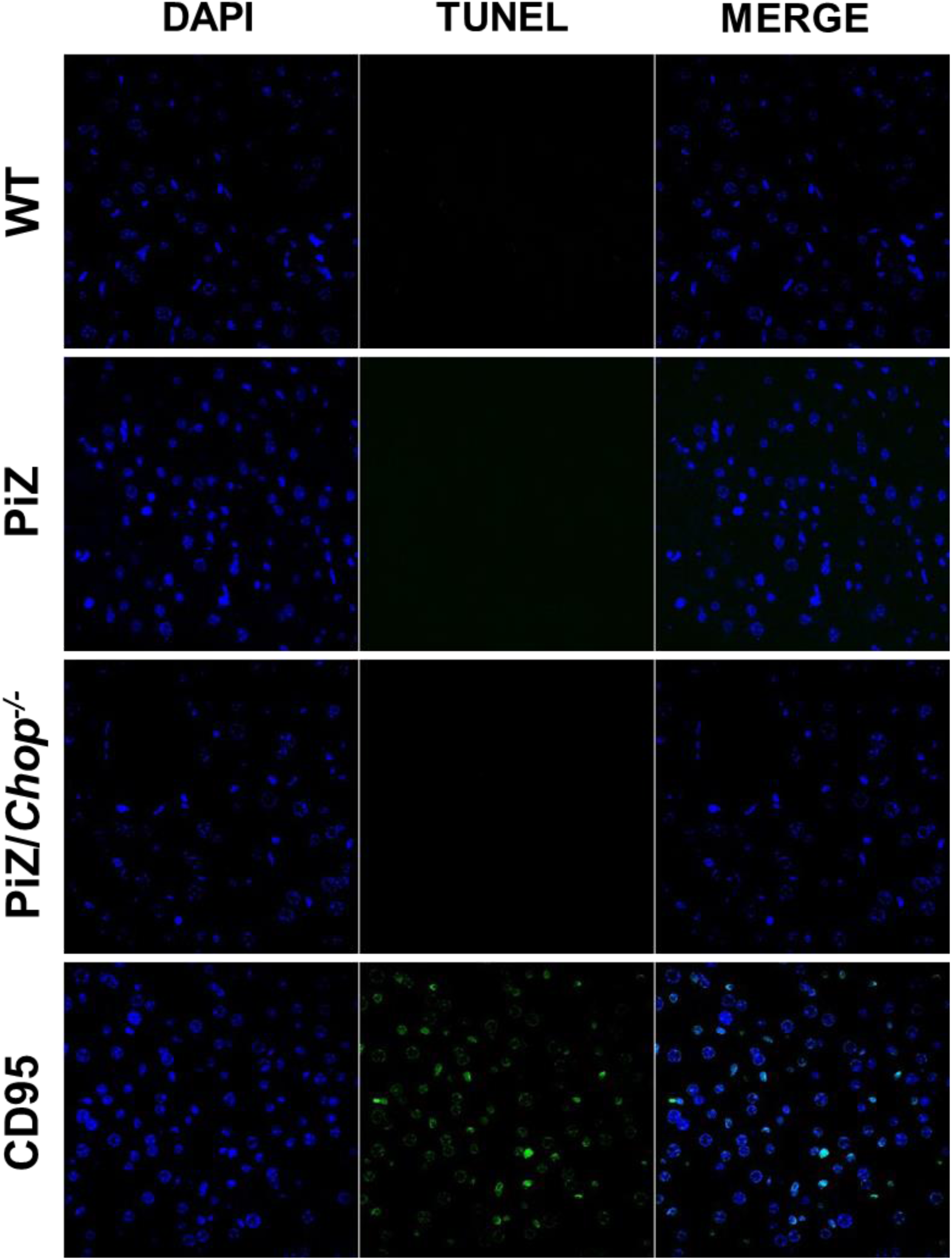
Representative TUNEL staining in 6-week old wild-type, PiZ and PiZ/*Chop*^*-/-*^ mouse livers (63X magnifications, n=3 per group). Livers of wild-type mice treated with 0.5 μg/g of body weight of CD95-activating antibody 6 hours prior tissue harvesting were used as positive control of the staining. *Abbreviations: WT, wild type.*

**Supplementary Figure S9:**
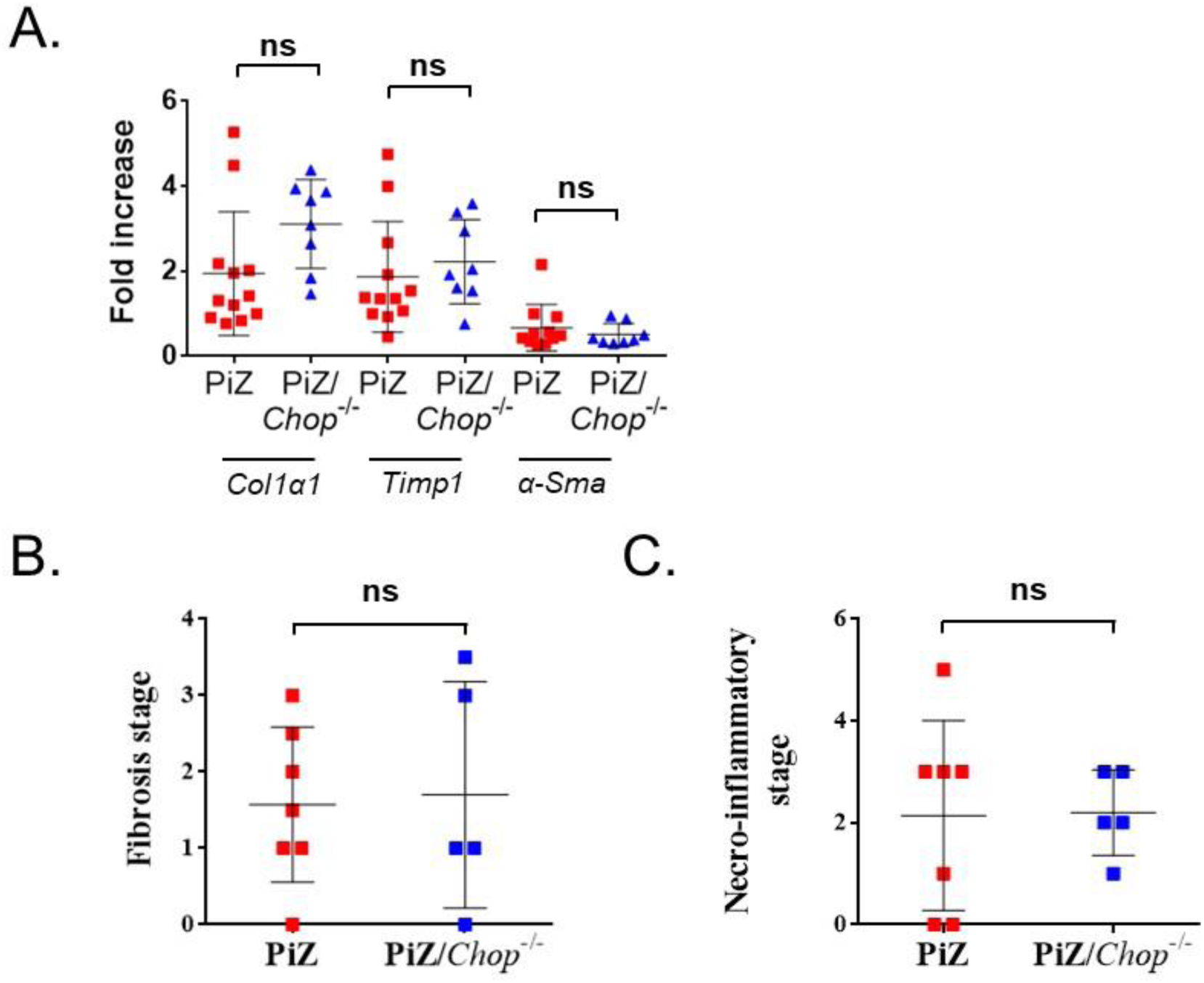
(**A**) Expression of genes associated to hepatic fibrosis (*Col1α1, Timp1, α-Sma*) in PiZ and PiZ/*Chop*^-/-^ mice at 69 weeks of age. *β2-Microglobulin* was used for normalization. (**B**, **C**) METAVIR score on liver sections from 69-weeks old PiZ and PiZ/*Chop*^-/-^ mice. Averages ± standard deviations are shown. Abbreviations: ns, not statistically significant.

**Supplementary Figure S10:**
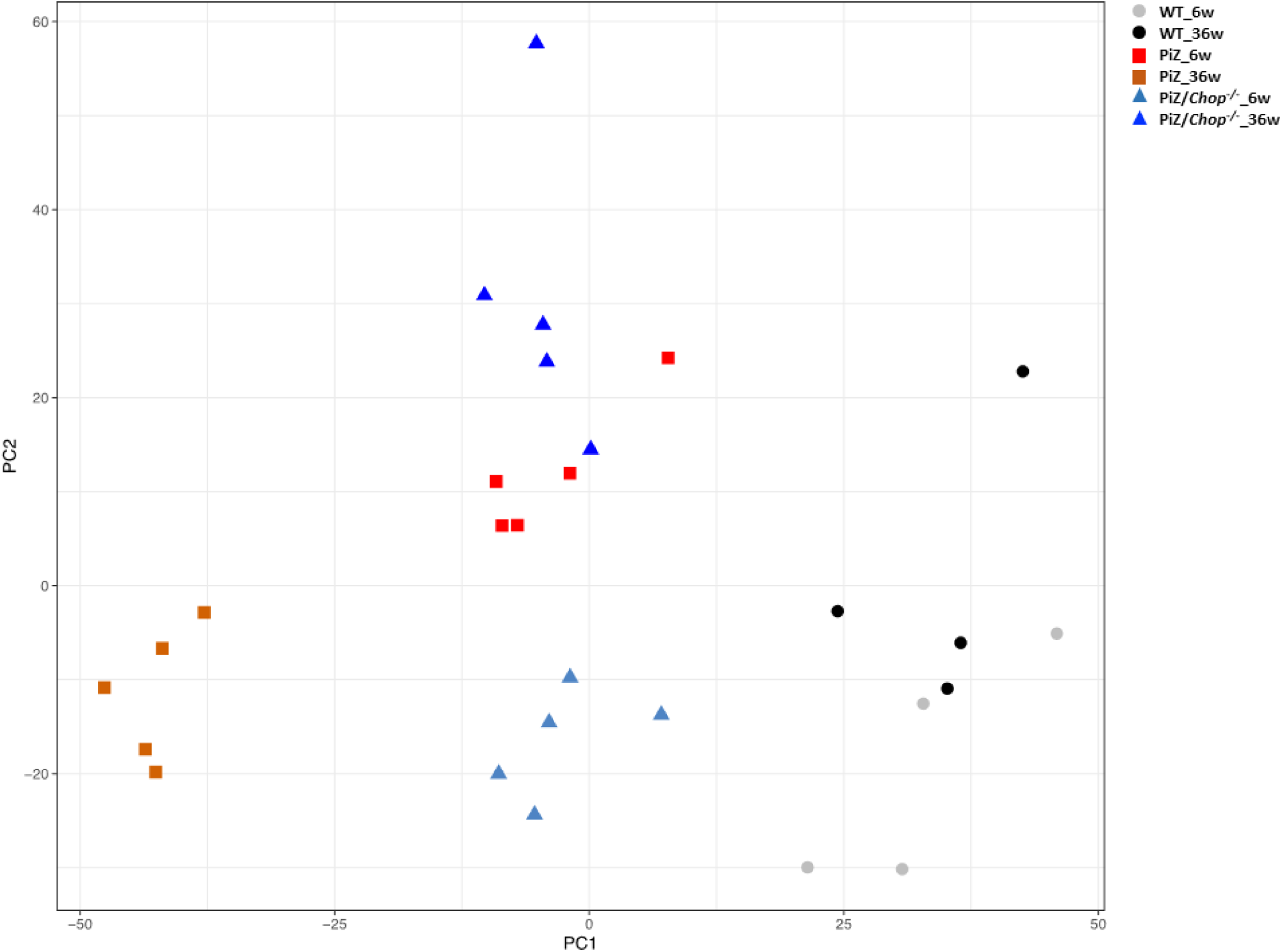
Principal component analysis (PCA) showing the distribution of gene expression profiles of wild-type, PiZ, and PiZ/*Chop*^-/-^ mouse livers at 6- and 36-weeks of age. *Abbreviations: WT, wild-type; 6w, 6 weeks; 36w, 36-weeks of age.*

**Supplementary Figure S11:**
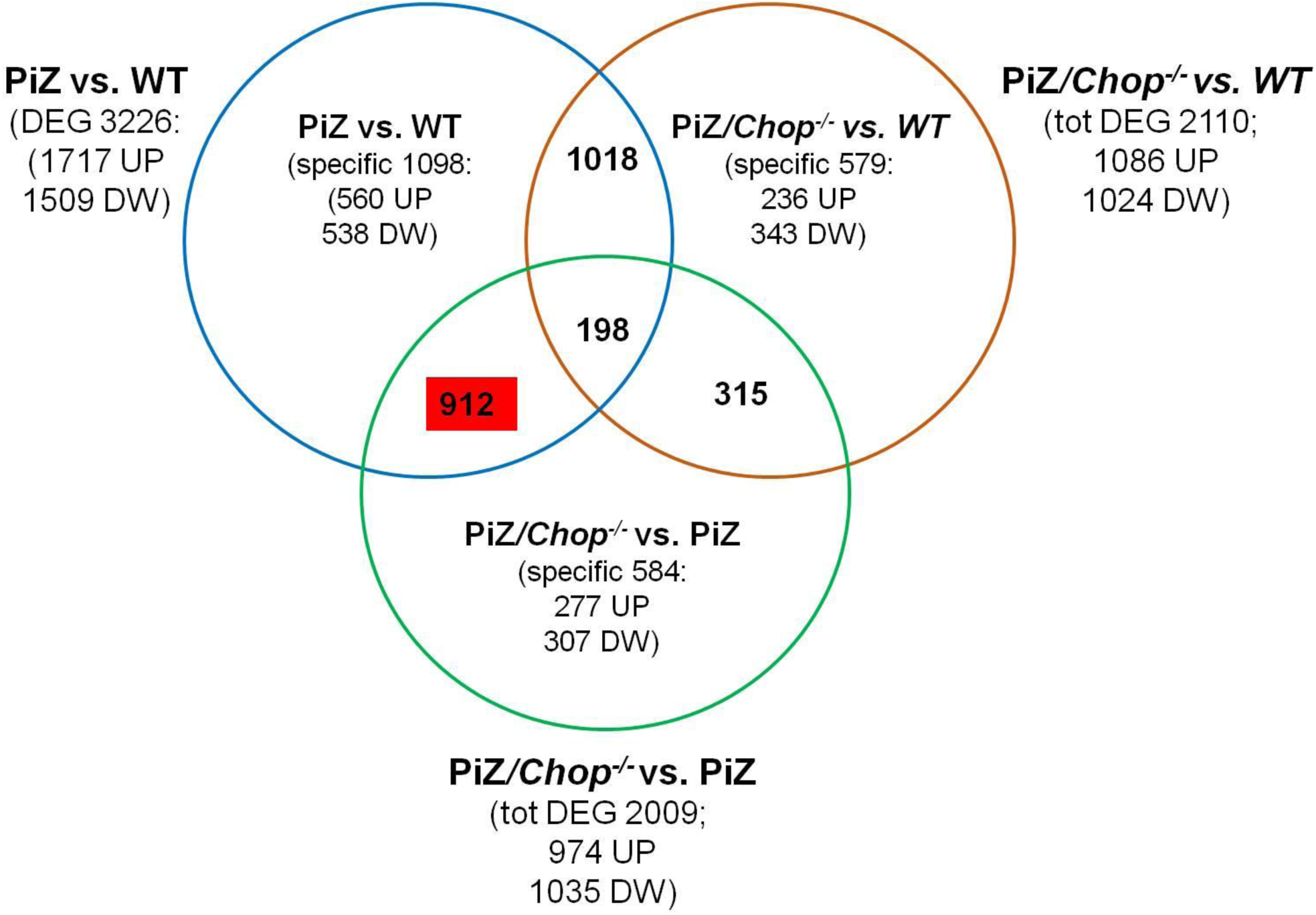
VENN diagram comparing the gene datasets of PiZ *vs.* wild-type, PiZ/*Chop*^*-/-*^ *vs.* wild-type, and PiZ/*Chop*^*-/-*^ *vs.* PiZ at 6-weeks of age. *Abbreviations: DEG, differentially expressed genes; DW, downregulated; UP, upregulated; WT, wild-type.*

**Supplementary Figure S12:**
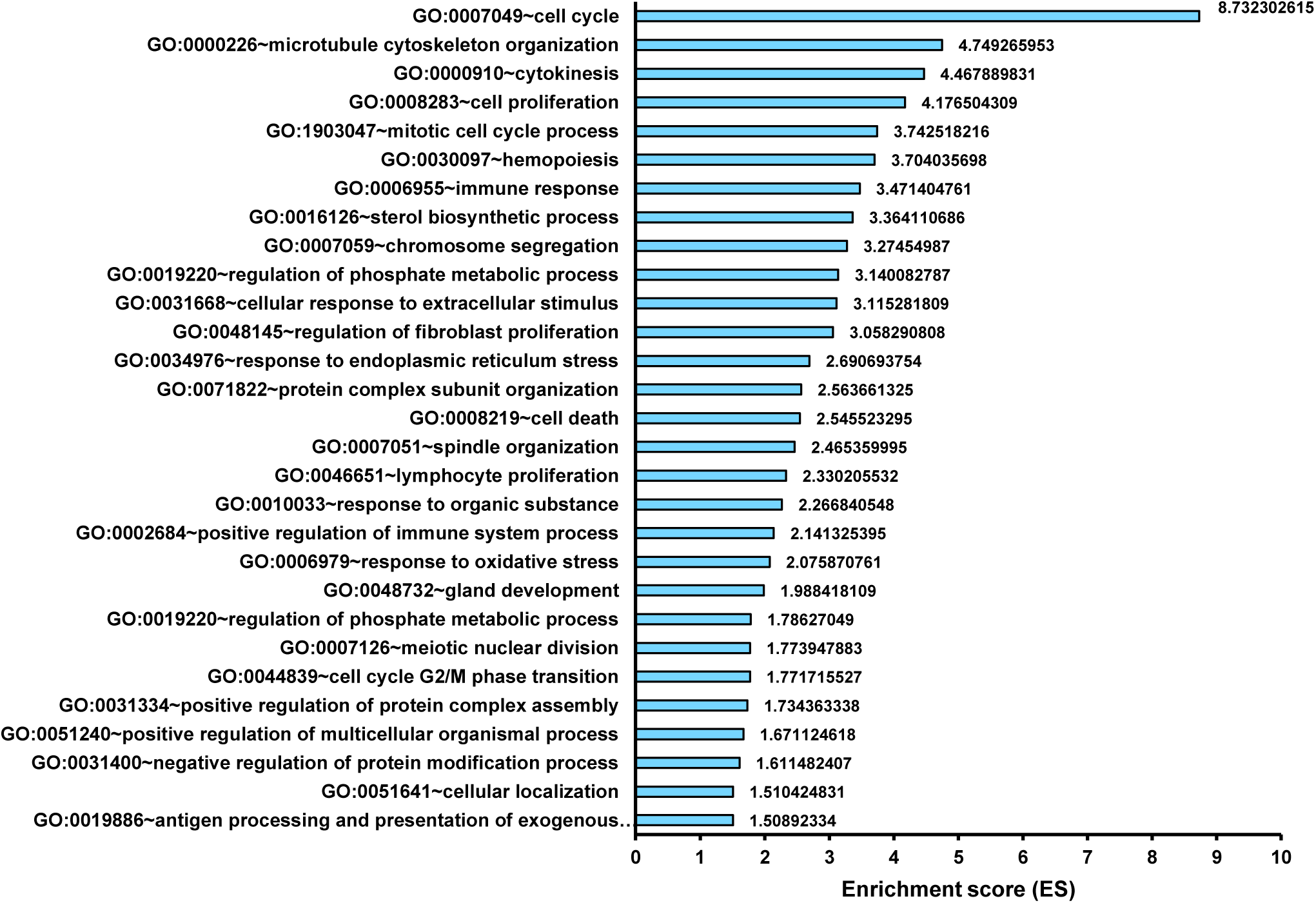
Functional annotation clustering analysis on differentially expressed genes in PiZ/*Chop*^-/-^ *vs.* PiZ livers at 6 weeks of age. Significantly downregulated clusters (enrichment score >1.5) are shown. Clusters are named after their most significant GO term. Complete lists of GO term forming the clusters are available in **Supplementary data**.

**Supplementary Figure S13:**
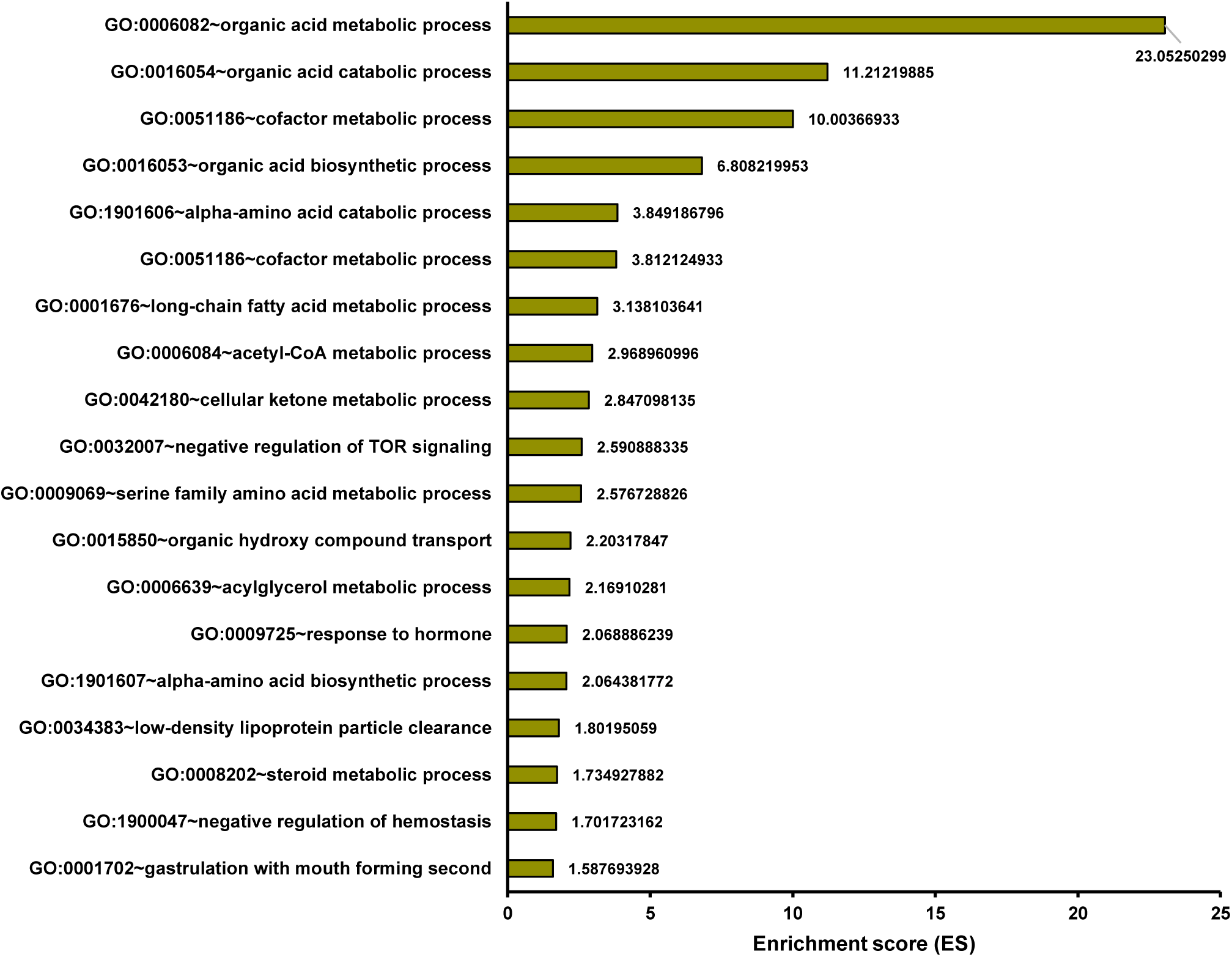
Functional annotation clustering analysis on differentially expressed genes in PiZ/*Chop*^-/-^ *vs.* PiZ livers at 6 weeks of age. Significantly upregulated clusters (enrichment score >1.5) are shown. Clusters are named after their most significant GO term. Complete lists of GO term forming the clusters are available in **Supplementary data**.

**Supplementary Figure S14:**
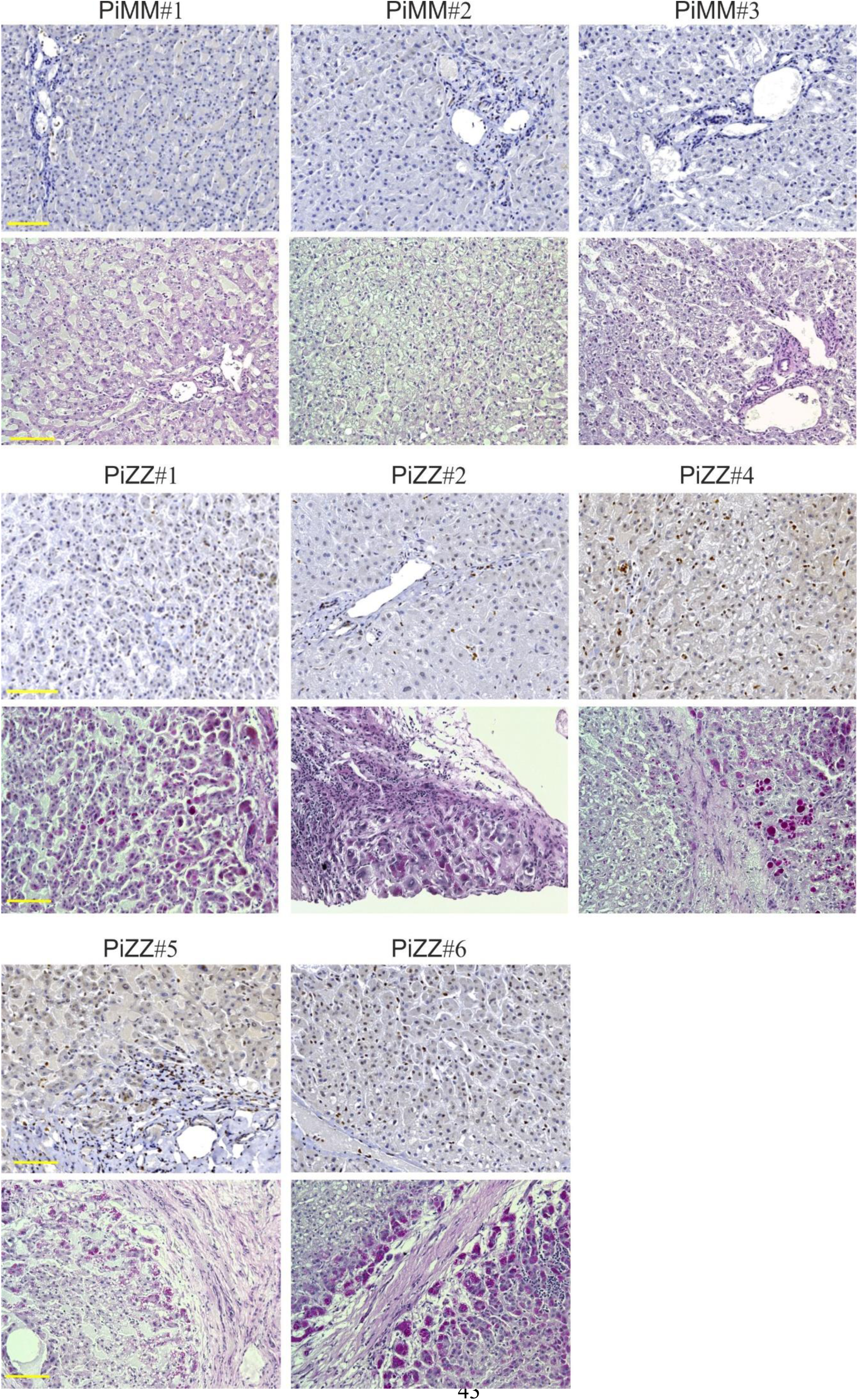
Immunohistochemistry with anti-CHOP antibody (upper panels) and corresponding PAS-D staining (lower panels) in additional livers of PiZZ patients with end-stage liver disease who underwent liver transplantation (see also **Supplementary table S5**) and age-matched PiMM controls who also underwent liver transplantation as controls (20X magnifications; scale bar, 100µm).

## Supplementary Methods

### Isolation of parenchymal and non-parenchymal cells

Parenchymal and non-parenchymal cells were isolated from livers of 6-week old PiZ mice by a modified protocol based on pronase/collagenase digestion (44). In brief, mouse livers were perfused through the inferior vena cava with EGTA solution followed by enzymatic digestion with pronase (Sigma-Aldrich P5147) and then collagenase type D (Roche 11088882001). Next, livers were harvested, and liver cells were disassociated by digestion with pronase/collagenase solution and filtered through a nylon filter (Corning 431751) to remove undigested tissues and debris. The resulting cell suspension was centrifuged at 70g for 3 minutes at 4°C. Supernatant containing non-parenchymal cells was collected. The cell pellet was washed 3 times with William’s medium E (Gibco 22551-022) and centrifuged at 70g for 3 minutes at 4°C to obtained hepatocytes. The non-parenchymal liver cell fraction was obtained after the centrifugation at 600g for 10 minutes at 4°C. The cell pellet was washed with HBSS without Ca^2+^ and Mg^2+^ 1x (Gibco 14175-095) and centrifuged at 70g for 3 minutes at 4°C to minimize hepatocyte contamination. The final cell suspension was centrifuged at 600g for 10 minutes at 4°C to obtain non-parenchymal cell fraction. Real time PCR for albumin gene was performed to evaluate enrichment for the parenchymal fraction and gene expression was normalized to 18S ribosomal RNA.

### ATZ Immunoblot

Preparation of soluble and insoluble fractions from livers was performed according to previous studies (45) and immunoblotting was performed using antibodies against total AAT and ATZ polymer (**Table S2**).

### Tissue stainings

Immunofluorescence staining of apoptotic cells was performed by terminal deoxynucleotidyl transferase-mediated deoxyuridine triphosphate nick-end labeling (TUNEL) using In Situ Cell Death Detection Kit (Roche Molecular Biochemicals) according to the manufacturer’s instructions. Mice injected with 0.5 μg/g of body weight of CD95-activating antibody (BD Biosciences) for 6 hours were used as positive control. The image capture was performed using Leica DM5000 microscope.

The mean area and number of the ATZ globules by liver PAS-D staining were quantified using the ImageJ plug-in “Analyze particles” on the segmented images. Five different fields of view selected randomly for n=3 PiZ and PiZ/*Chop*^*-/-*^ mice were used for the quantification. The image capture was performed using Leica DM5000 microscope.

Sirius red staining was performed on 5-μm liver sections rehydrated and stained for 1 hour in 0.1% Sirius red (Sigma) in saturated aqueous solution of picric acid. After two changes of acidified water (0.5% acetic acid in water), the sections were dehydrated, cleared in xylene and mounted in a resinous medium. Image capture was performed using the Leica DM5000 microscope. METAVIR scoring system (46) was used to determine the severity of hepatic fibrosis and necro-inflammatory stage. All samples were evaluated by the same pathologist (S.C.) who was blinded about the laboratory data.

---

**Supplementary Table S1:** Pathology features PiZZ and PiMM controls.

|  | **METAVIR** |  | **PAS-D** |
| --- | --- | --- | --- |
|  | **Fibrosis** | **Necro-inflammation** |  |
| PiZZ #1 | 4 | 5 | +++ |
| PiZZ #2 | 3 | 3 | + |
| PiZZ #3 | 4 | 5 | ++ |
| PiZZ #4 | 4 | 2 | +++ |
| PiZZ #5 | 4 | 4 | ++ |
| PiZZ #6 | 4 | 4 | +++ |
| PiMM #1 | 1 | 1 | - |
| PiMM #2 | 1 | 1 | - |
| PiMM #3 | 1 | 1 | - |

PAS-D score: +, focal periportal/peri-septal deposits; ++, multifocal periportal/peri-septal deposits; +++, diffuse multifocal periportal/peri-septal deposits or multifocal periportal/peri-septal and centrilobular.

**Supplementary Table S2.** Clinical and pathology features PiZZ and control adults and children.

| ID | Gender | Age (years) | Diagnosis and clinical features |
| --- | --- | --- | --- |
| **Healthy controls** |  |  |  |
| Adults |  |  |  |
| H2 | M | 63 | - |
| H3 | M | 58 | - |
| H4 | M | 55 | - |
| H6 | M | 53 | - |
| H7 | M | 50 | - |
| H9 | M | 25 | - |
| Children |  |  |  |
| H21 | M | 5 | - |
| H22 | M | 4 | - |
| H23 | F | 3.5 | - |
| H24 | M | 2 | - |
| H25 | F | 1.5 | - |
| H11 | M | 12 | - |
| H12 | M | 11 | - |
| H13 | F | 10 | - |
| H14 | M | 9 | - |
| H15 | F | 9 | - |
| H16 | F | 8 | - |
| H18 | F | 6 | - |
| **PiZZ** |  |  |  |
| Adults |  |  |  |
| A2 | M | 63 | ZZ |
| A3 | M | 58 | *ZZ* |
| A4 | M | 55 | *ZZ* |
| A6 | M | 53 | *ZZ* |
| A7 | M | 50 | *ZZ* |
| A9 | M | 26 | *ZZ* |
| Children |  |  |  |
| A21 | M | 4.8 | *ZZ* |
| A22 | M | 3.8 | *ZZ* |
| A23 | F | 3.5 | *ZZ* |
| A24 | M | 1.7 | *ZZ* |
| A25 | M | 1.3 | *ZZ* |
| A26 | M | 1.6 | *ZZ* |
| A27 | M | 11 months | *ZZ* |
| A28 | M | 10 months | *ZZ* |
| A29 | F | 7 months | *ZZ* |
| A30 | F | 7 months | *ZZ* |
| A11 | M | 12 | *ZZ* |
| A12 | M | 11 | *ZZ* |
| A13 | F | 10 | *ZZ* |
| A14 | M | 9.6 | *ZZ* |
| A15 | F | 8.8 | *ZZ* |
| A16 | F | 8.4 | *ZZ* |
| A17 | F | 8.4 | *ZZ* |
| A18 | F | 6.3 | *ZZ* |
| Diseased controls |  |  |  |
| Adults |  |  |  |
| D2 | M | 61 | alcoholic cirrohis, HBV- HCV- |
| D3 | M | 58 | cirrhosis, HBV- HCV- |
| D4 | M | 54 | cirrhosis, HBV- HCV- |
| D6 | M | 55 | cirrhosis |
| D7 | M | 50 | cirrhosis, HCV- |
| D9 | M | 25 | cirrhosis |
| Children |  |  |  |
| D21 | M | 3.4 | biliary atresia or cirrhosis |
| D23 | F | 2.3 | biliary atresia or cirrhosis |
| D24 | M | 1.6 | biliary atresia or cirrhosis |
| D25 | M | 1.2 | biliary atresia or cirrhosis |
| D26 | M | 1.5 | biliary atresia or cirrhosis |
| D12 | M | 13.0 | biliary atresia or cirrhosis |
| D13 | F | 10.3 | biliary atresia or cirrhosis |
| D14 | M | 8.8 | biliary atresia or cirrhosis |
| D15 | F | 8.8 | biliary atresia or cirrhosis |
| D18 | F | 7.8 | biliary atresia or cirrhosis |

**Supplementary Table S3:**
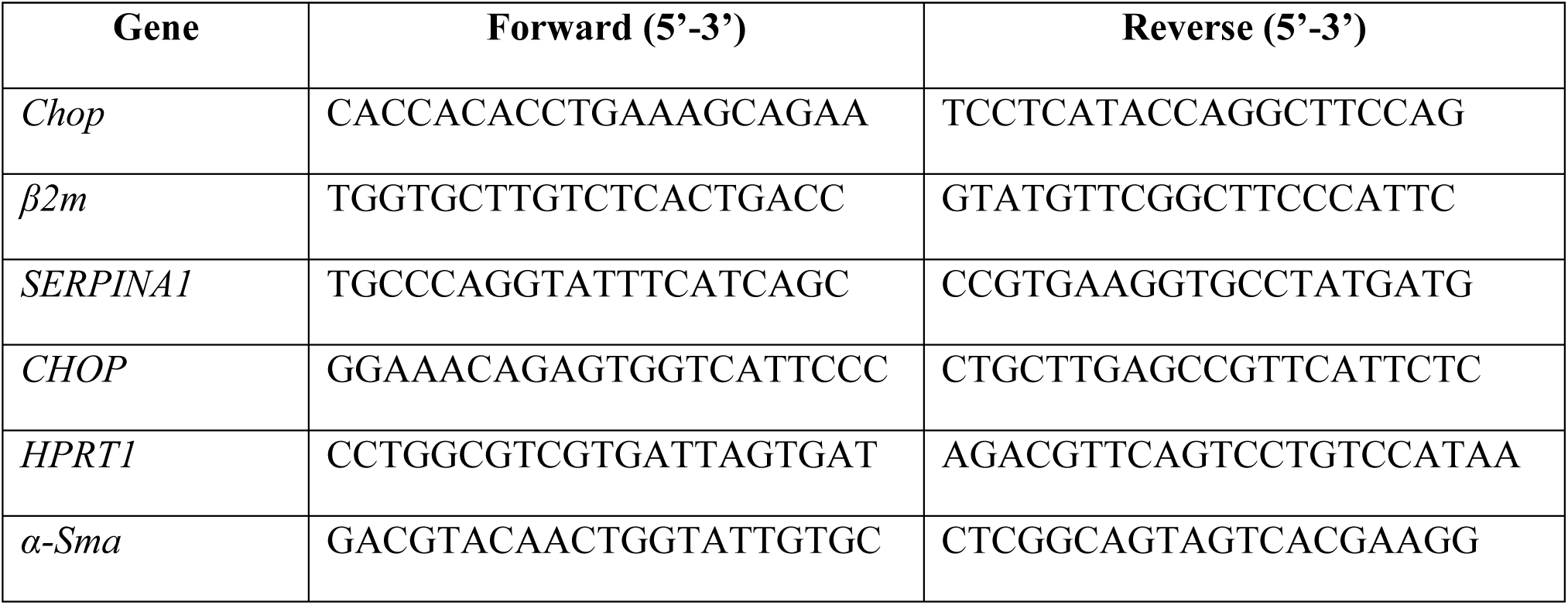
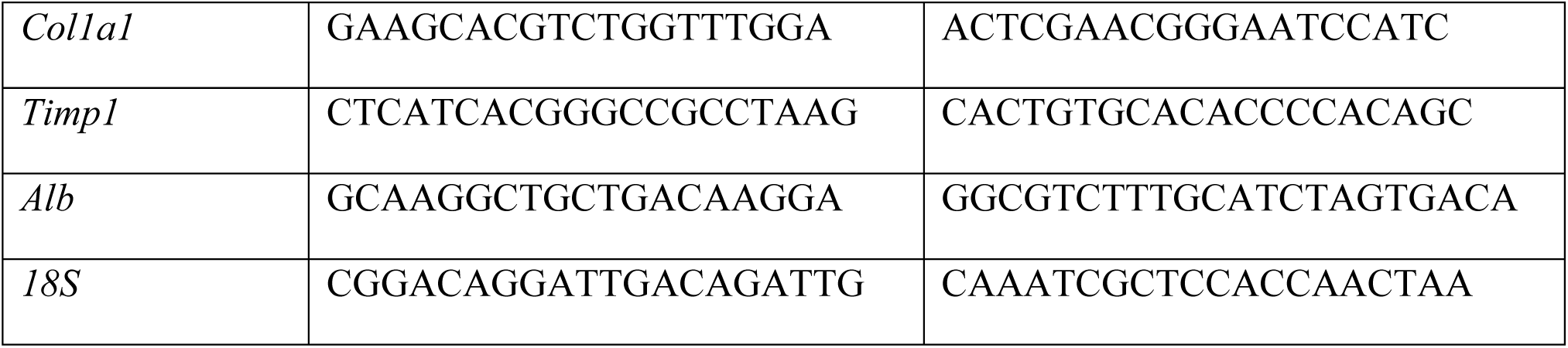
Sequence of primers used for qPCR.

**Supplementary Table S4:** Primary antibodies and dilutions.

| **Antibody** | **Manufacturer** | **WB**<br>**dilution** | **IHC**<br>**dilution** | **IF**<br>**dilution** | **IP**<br>**dilution** | **ELISA** | **ChIP** |
| --- | --- | --- | --- | --- | --- | --- | --- |
| anti-CHOP | Abcam<br>(ab11419) | 1/500 | 1/100 | - | - | - | - |
| anti-CHOP | LifeSpan<br>Biosciences<br>(LS-B3652) | - | 1/40 | - | - | - | - |
| anti-GAPDH | Santa Cruz<br>Biotech<br>(sc-365062) | 1/3,000 | - | - | - | - | - |
| anti-LAMIN-A | Abcam<br>(ab26300) | 1/1,000 | - | - | - | - | - |
| anti-AAT | DAKO<br>(A0012) | - | - | - | - | 1/4,000 | - |
| anti-ATZ<br>polymer | Hycult Biotech<br>(HM2289) | - | - | 1/100 | - | - | - |
| anti-FLAG | Sigma<br>(F1804) | - | - | - | 1/200<br>(3 µg) | - | - |
| anti-FLAG | Sigma<br>(F7425) | 1/2,000 | - | - | - | - | - |
| anti-CALNEXIN | Enzo Life<br>Sciences<br>(ADI-SPA-860) | 1/1,000 | - | - | - | - | - |
| Anti-CHOP | Cell signaling<br>(2895) | - | - | - | - | - | 1/100 |
| Anti-Myc | Cell signaling<br>(2278) | 1/1,000 | - | - | - | - | - |
| Anti-phospho-JUN | Cell signaling<br>(3270) | 1/1,000 | - | - | - | - | 1/100 |
| IgG mouse | Santa Cruz<br>Biotech<br>(sc-2025) | - | - | - | 1/80<br>(3 µg) | - | - |

*Abbreviations: WB, western blot; IHC, immunohistochemistry; IF, immunofluorescence; IP, immunoprecipitation; ELISA, enzyme-linked immunosorbent assay; ChIP, chromatin immunoprecipitation.*

**Supplementary Table S5:**
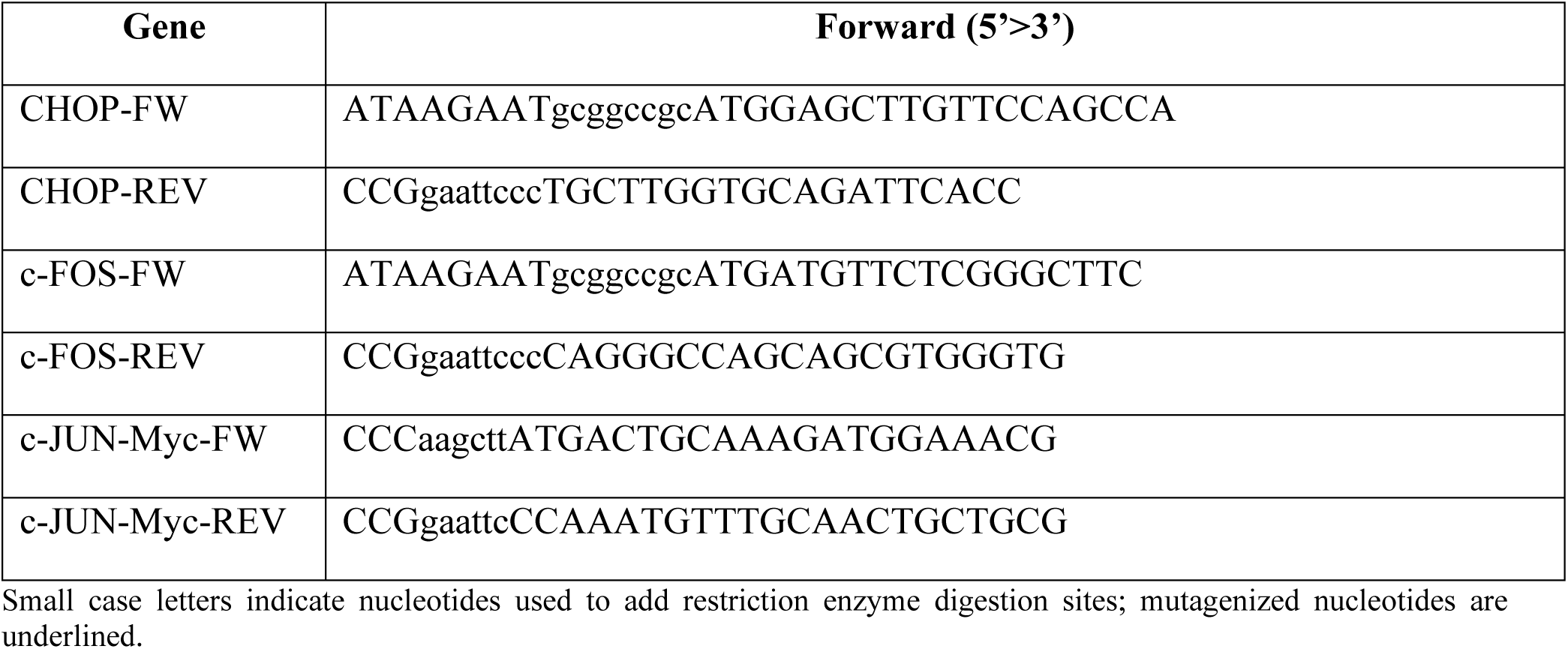
Forward (FW) and reverse (REV) primers for cloning of transcription factors into plasmids used in luciferase assay and immunoprecipitations.

**Supplementary Table S6:**
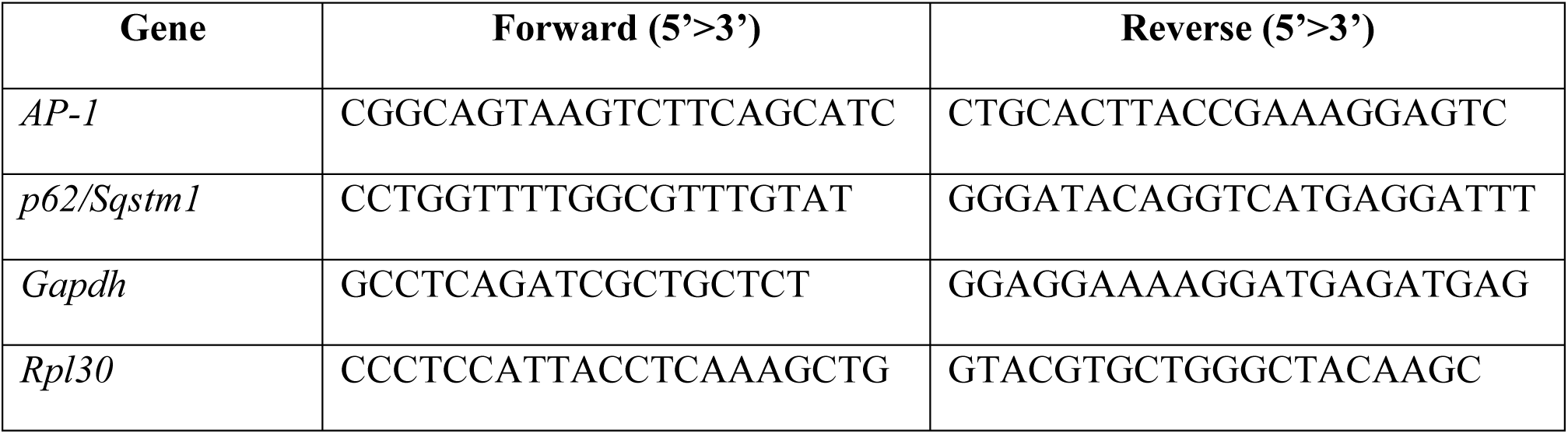
qPCR primers for chromatin immunoprecipitation (ChIP).

